# Lateral occipitotemporal cortex encodes perceptual components of social actions rather than abstract representations of sociality

**DOI:** 10.1101/722249

**Authors:** Moritz. F. Wurm, Alfonso Caramazza

**Affiliations:** Center for Mind/Brain Sciences, University of Trento, 38068 Rovereto (TN), Italy; Cognitive Neuropsychology Laboratory, Harvard University, Cambridge, MA 02138 USA

## Abstract

Neuroimaging studies suggest that areas in the lateral occipitotemporal cortex (LOTC) play an important role in the perception of social actions. However, it is unclear what precisely about social actions these areas represent: perceptual features that may be indicative of social actions – such as the presence of persons in a scene, their orientation toward each other, and in particular the directedness of action movements toward persons or other targets – or more abstract representations that capture whether an action is meant to be social. In two fMRI experiments, we used representational similarity analysis (RSA) to test whether LOTC is sensitive to perceptual action components important for social interpretation and/or more general representations of sociality (Experiment 1) and implied person-directedness (Experiment 2). We found that LOTC is sensitive to perceptual action components (person presence, person orientation, and action directedness toward different types of recipients). By contrast, more general levels of sociality and implied person-directedness were not captured by LOTC. Our findings suggest that regions in LOTC provide the perceptual basis for social action interpretation but challenge accounts that posit specialization at more general levels sensitive to social actions and sociality as such. We propose that the interpretation of an action – in terms of sociality or other intentional aspects – arises from the interaction of multiple areas in processing relevant action components in a situation-dependent manner.

## 1. Introduction

A current question in neuroscience concerns how knowledge about different types of actions, such as locomotion, object manipulation, or communication actions, is organized in the brain (Abdollahi et al., 2013; Handjaras et al., 2015; Tarhan and Konkle, 2017; Tucciarelli et al., 2019; Wurm and Caramazza, 2019; Wurm et al., 2017). A class of actions that has received particular interest are social actions (for a review see Quadflieg and Koldewyn, 2017), which can be defined as actions that take into account the behavior of others and are thereby oriented in their course (Weber, 1978). Social actions may have particular relevance for observers, as they can indicate an attack or approach (at a group member or the observing person herself) and are informative for understanding social relationships (e.g., who is friend or foe of whom?) and personality traits (e.g., is she a generous person?). It therefore appears plausible that specialized neural mechanisms evolved to process social actions. In potential support of this view, neuroimaging studies found that observing videos or point-light displays of interacting (e.g. *dancing*, *communicating*) persons vs. two non-interacting persons (Centelles et al., 2011; Isik et al., 2017; Pierno et al., 2008; Walbrin et al., 2018) or videos or pictures of incongruent vs. congruent interactions between persons (Petrini et al., 2014; Quadflieg et al., 2015) increases neural activity in brain regions in proximity to the visual system such as the posterior superior temporal sulcus (pSTS) and lateral occipitotemporal cortex (LOTC) as well as more anterior/dorsal regions such as lateral prefrontal and inferior parietal areas.

What precisely about social actions could be captured in these regions? One possibility is that these regions are specialized for perceptual features that are predictive of social actions, such as the presence of persons in a common place (Isik et al., 2017; Wagner et al., 2011; Walbrin et al., 2018) and the orientation of persons toward each other (Centelles et al., 2011; Isik et al., 2017; Kujala et al., 2012; Walbrin et al., 2018). These features, however, are neither necessary nor sufficient for social interpretation. For example, persons might face each other without interacting, and persons can interact without facing each other and even without being present at the same place (e.g. *communicating via phone*). Thus, early stages of processing would be specialized not for social actions as such but rather for salient features that are important for interpreting an action as being social. From a theoretical perspective, it is generally questionable whether there are brain regions that selectively represent social actions but not other kinds of actions. This view would presuppose a categorical distinction into social and nonsocial actions. However, the transition from social to nonsocial action appears to be continuous rather than categorical (e.g. from *greeting someone* to *preparing a dinner* to *reading a book*), and whether an action is perceived as social often depends on subtle contextual cues (e.g. *writing a letter* vs. *a to-do list*). Also, for certain subtypes of social actions, such as social interactions in which more than one person are actively involved, a segregation is often unclear and context-dependent (e.g. two persons *taking a walk together* vs. *coincidentally walking next to each other*). Another possibility is therefore that specialization occurs at a higher, more abstract level that is independent of salient perceptual features. Following this view, this level of representation might integrate different action-related aspects (e.g. body movements as well as other contextual cues about objects, persons, and the specific situation) that together specify the interpretation of whether or not an action is implied to be social, i.e., takes into account the behavior of others and is thereby oriented in its course (sociality hereafter).

We recently found that bilateral LOTC discriminated actions judged as being social (e.g. *giving an object to another person*, *making an agreement gesture to another person*) from non-social actions (e.g. *opening an object*, *scratching the arm*) (Wurm et al., 2017). Notably, the representations sensitive to this distinction generalized across different, perceptually heterogeneous actions (e.g. across *giving* and *making an agreement gesture*), suggesting that these representations captured more general aspects that are independent of action-specific details, such as specific body movements or the specific meaning of an action (e.g. the physical transfer of an object to another person vs. informing another person about agreement). Additionally, in both social and nonsocial actions, two persons were present that were facing each other, ruling out basic perceptual explanations. The findings therefore are consistent with the hypothesis that LOTC encodes the sociality of actions at an abstract, perceptually invariant level of representation. However, the social actions involved a movement made in the direction of the other person, whereas the non-social actions were directed toward other targets (inanimate objects or the actress’ own body). Hence, it is possible that the distinction between social and non-social actions was driven by the directedness of the actions toward persons vs. other targets. Notably, besides the presence of persons in a scene and their orientation toward each other, the directedness of actions is a particularly salient cue indicating whether or not an action could be social. Therefore, neural specialization to detect action directedness might be evolutionary advantageous. To date, however, it remains unexplored whether there are brain regions specifically tuned to action directedness.

Taken together, there is substantial evidence that areas around LOTC and pSTS play an important role in the perception of social actions. However, what precisely these regions encode – an integrational level that captures sociality or rather certain “protosocial” components that may be indicative of sociality, such as the presence of persons in a scene, the orientation of persons toward each other and/or the directedness of action movements into the direction of others – remains controversial.

Here, we scrutinize the role of LOTC in representing sociality and more basic aspects of observed actions, with a particular focus on action directedness, under carefully controlled testing conditions. Using fMRI-based representational similarity analysis (RSA) (Kriegeskorte et al., 2008) we tested for different levels of representation associated with the observation of simple actions – moving an object from X to Y – that vary in terms of being interpreted as being social (Figs. 1 and 2). Notably, we did not use complex scenarios, e.g. those in which more than one person is moving, to avoid effects on cognitive processes not directly related to action recognition and interpretation. For example, previous studies contrasted social interactions between persons with two independent (potentially conflicting, and often less complex) actions, e.g., gesturing to sit down followed by the other person sitting down vs. a rowing and a sawing person next to each other (Centelles et al., 2011; Isik et al., 2017; Petrini et al., 2014; Pierno et al., 2008; Quadflieg et al., 2015; Walbrin et al., 2018). Activation differences therefore might be due to differences in complexity or other processes, such as detecting a semantic match or mismatch between two persons’ actions. Thus, our experimental paradigm focused on the minimal conditions that are sufficient for an action to be perceived as social. Notably, behavioral evidence demonstrated that mutual interaction is not necessary for the perception of social aspects in comparable “giving” and “taking” actions, even in absence of tasks that explicitly require attention to social dimensions (Tatone, Geraci, & Csibra, 2015). Since we aimed at investigating automatic recognition processes, we only ensured the participants’ attentiveness to the actions by using a catch trial task (actions that deviated along a non-social dimension from the experimental trials). Experiment 1 focused on representations of socially relevant cues (e.g. presence of other persons, directedness of actions toward persons vs. other targets, the orientation of persons relative to each other) vs. perception-invariant representations that capture the subjective interpretation whether an action is perceived as being social or not. This experiment revealed that LOTC is sensitive to perceptual action components – most importantly the directedness of actions toward different types of recipients – but not to sociality. Experiment 2 aimed at testing the level of generality of action direction representations in more detail: The presence of another person in a scene and action movements made in the direction of that person are salient cues important for interpreting an action as being social. However, actions can also be directed toward the observer, who is typically part of an action scene. Movements made in the direction of the observer could therefore represent similarly salient cues for social interpretation. Experiment 2 tested whether LOTC contains a general level of *implied* person- directedness that represents actions directed toward the observing participant in a similar manner as actions directed toward other persons. Again, we found that LOTC is sensitive to the directedness of actions but not to more general levels of implied person-directedness. Our results thus suggest that LOTC represents action components that are important for interpreting an action as being social, rather than the interpretational level itself that captures whether an action is meant to be person-directed and social.

**Figure 1.**
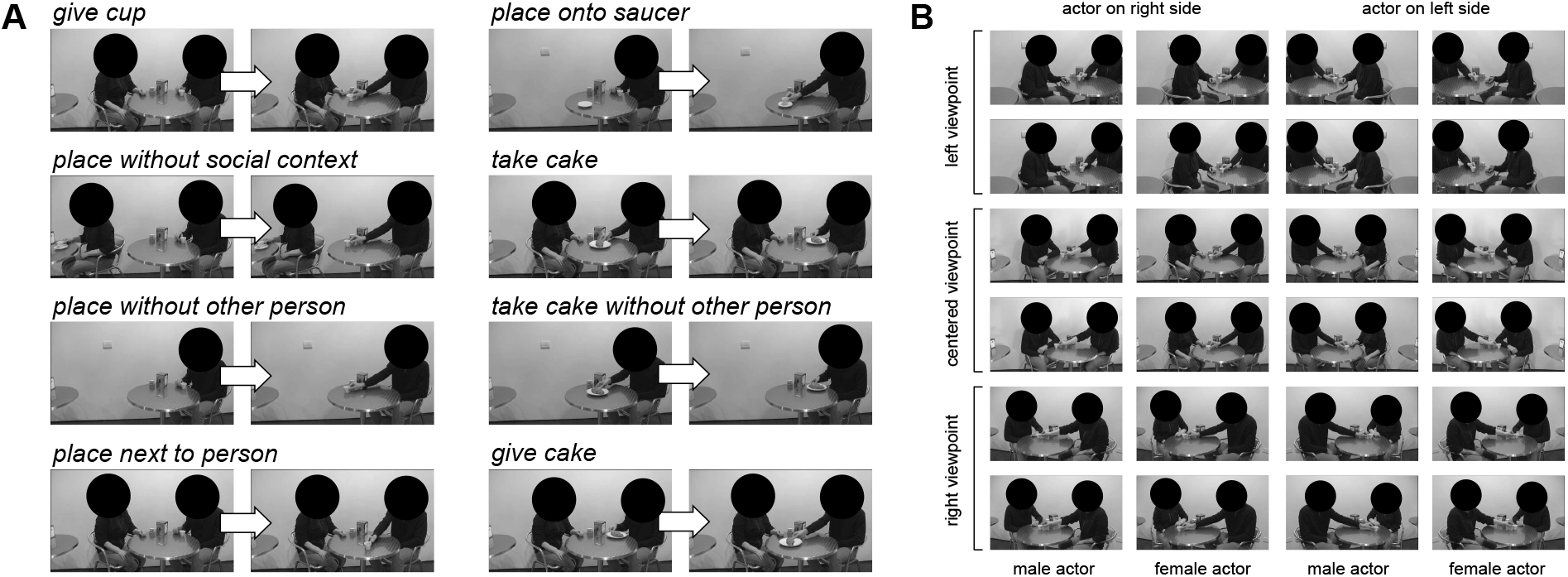
(A) Experimental design of Experiment 1. In eight video conditions (video duration = 2 s; first and middle frames of videos are shown), an actor moved an object into the direction of another person, toward an object (cup onto saucer), toward his- or herself, or not toward a specific entity, in the absence or presence of a third person facing or not facing the actor. Participants responded to occasionally presented catch trials in which meaningless variants of the actions were shown. (B) Stimulus variance (24 videos per condition). PLEASE NOTE THAT FACES WERE OBSCURED IN THE PREPRINT VERSION DUE TO POLICY CONSTRAINTS.

**Figure 2.**
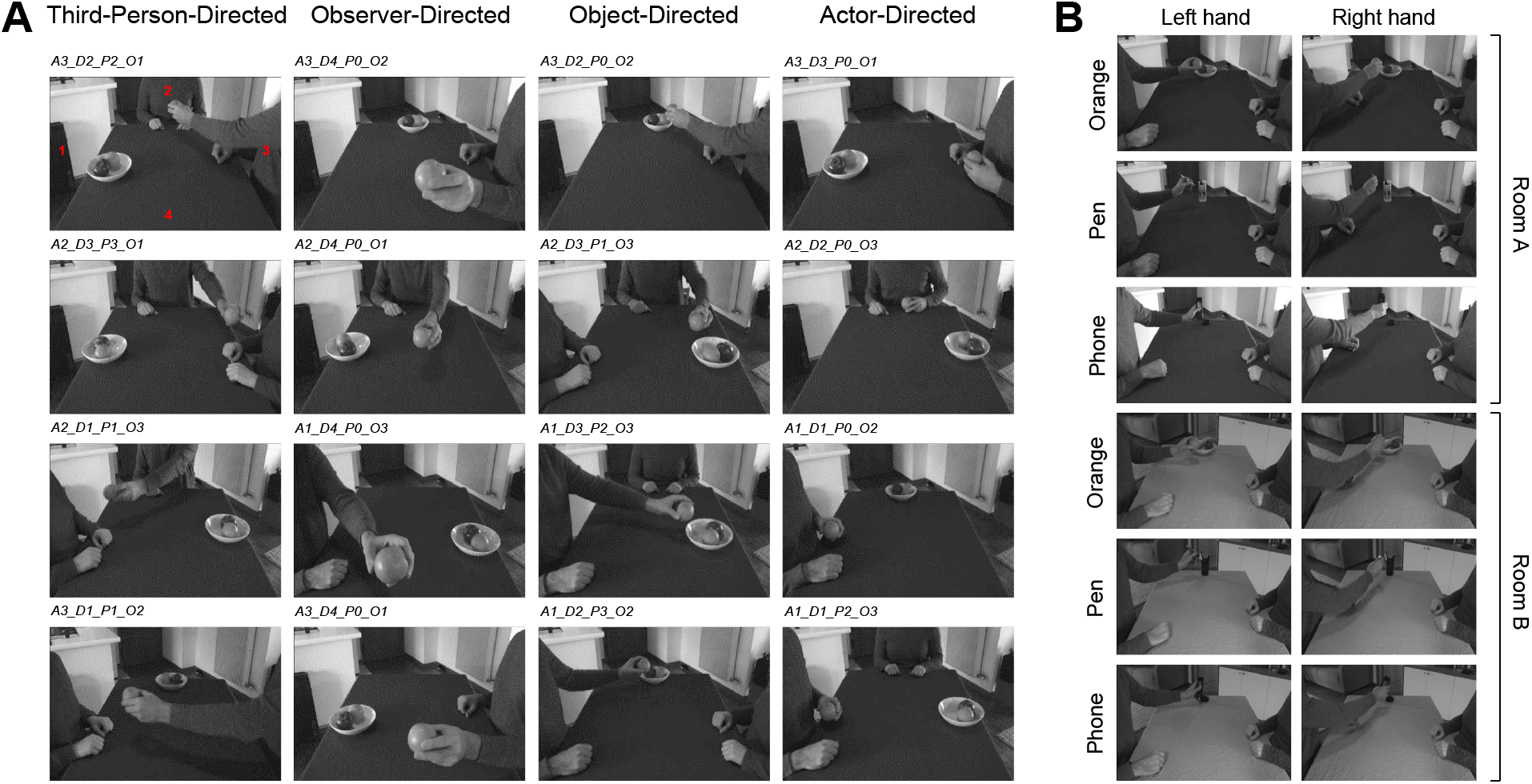
(A) Experimental design of Experiment 2. In 16 video conditions (video duration = 0.6 s; last frames of videos are shown), an actor moved an object toward another person, the observer, an object, or his- or herself, in the absence or presence of a third person. Locations of action targets (1,2,3,4; red numbers in upper left frame) and third person presence were varied to optimally minimize covariance between models of interest (shown in Fig. 3A; see Methods for details). Condition codes: A = actor position, D = direction of action (target location), P = third person position (0 for no third person present), O = object position. Participants responded to occasionally presented catch trials in which the action directions changed during the course of the trial. (B) Stimulus variance (12 videos per condition).

**Figure 3.**
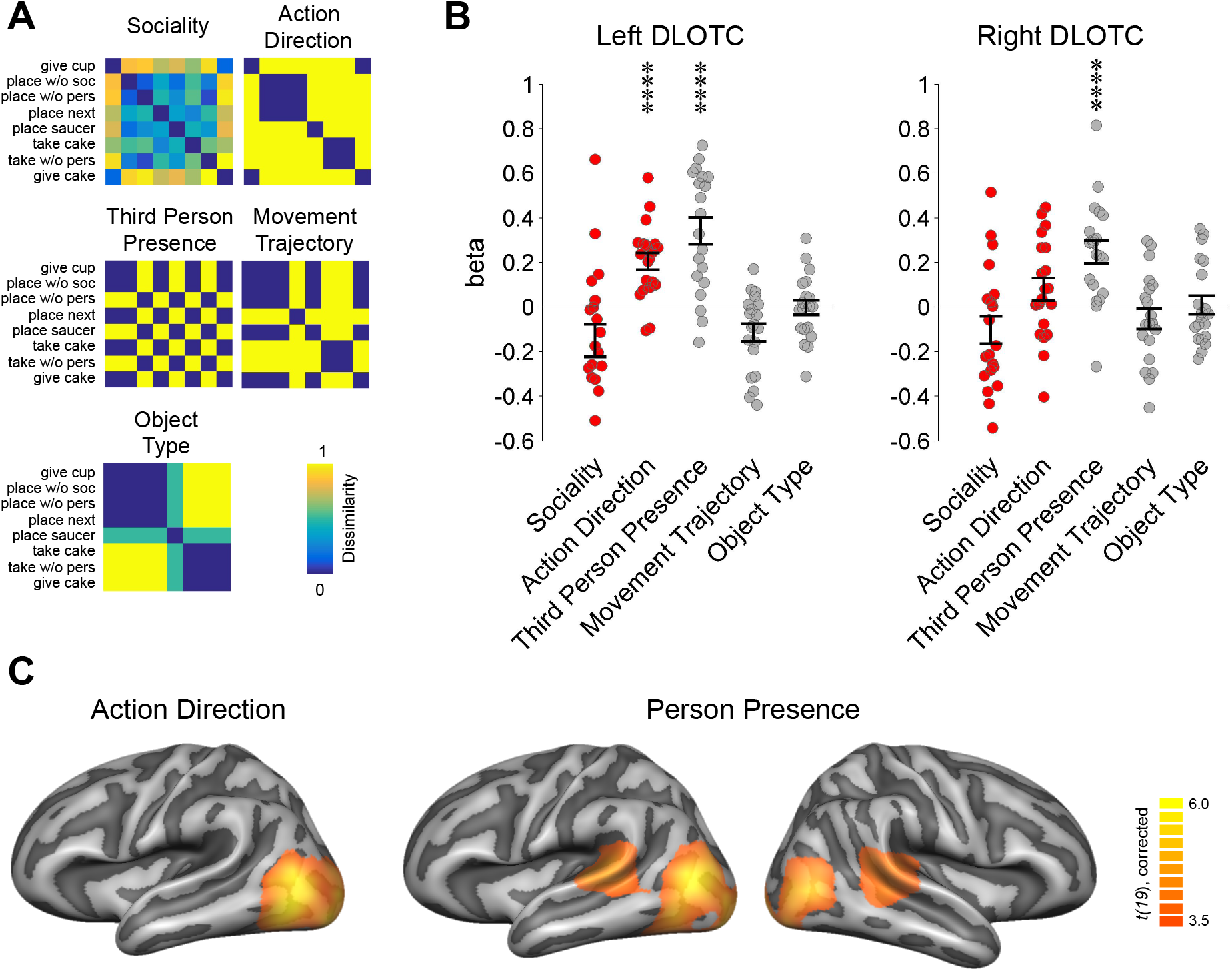
Multiple regression RSA targeting effects of sociality and action direction (Experiment 1). (A) Representational dissimilarity matrices (RDMs) of tested models. (B) ROI multiple regression RSA. ROIs (12 mm radius) were based on peak coordinates identified in the searchlight decoding for sociality (DLOTC) and transitivity (VLOTC) in Wurm et al. (2017). Dots indicate individual betas of the participants (main models of interest shown in red). Asterisks in indicate FDR-corrected (for number of models per test) effects: * p < 0.05, ** p > 0.01, *** p > 0.001, **** p > 0.0001. Error bars indicate SEM. (C) Searchlight multiple regression RSA using same RSA procedures as in the ROI analysis. Statistical maps are corrected for multiple comparisons (voxel threshold p = 0.001, corrected cluster threshold p = 0.05).

## 2. Material and Methods

### 2.1 Experiment 1

#### 2.1.1 Participants

Twenty healthy adults (13 females; mean age, 23.7 years; age range, 20-30 years) volunteered to participate in the experiment. All participants were right-handed with normal or corrected-to-normal vision and no history of neurological or psychiatric disease. Participants gave written informed consent prior to participation in the study. The experimental procedures were approved by the Ethics Committee for research involving human participants at the University of Trento, Italy.

#### 2.1.2 Stimuli

The stimulus set consisted of 2 s long videos showing 24 exemplars of eight actions (192 action videos in total, Fig. 1). The actions were filmed in a cafeteria setting. The videos showed either a single actor or an actor plus another passive person (factor Third-Person-Presence). The other person was sitting at either the same or a different table, thus facing or not facing the actor. Throughout the video, the other person did not make any arm, head, or eye movements. The actions consisted of moving an object toward the other person (*give cup, give cake, place without social context*), toward no specific entity, i.e., an unmarked location on the table (*place without another person*, *place next to person*), toward a target object (*place onto saucer*), or toward the actress or actor herself/himself (*take cake* and *take cake without another person*; factor Action Direction). This set of actions was chosen to manipulate the degree of sociality of the actions (factor Sociality) independently of Third-Person-Presence and Action Direction (see *Methods* section *Representational similarity matrices* for an analysis of putative correlations between the factors). Both persons looked at the action, i.e., the moving of the object, throughout the video.

To increase the stimulus variance, the actions were performed by two different actors filmed from three different perspectives (from the left, the right, and a centered camera viewpoint; Fig. 1B). For each perspective, each actor performed each action two times (resulting in six different action instantiations per actor). To further increase perceptual variance, each video was flipped, which resulted in additional action instantiations with the actor sitting either on the left or the right side of the table using either the left or the right hand for the actions.

Catch trials consisted of six exemplars of each of the eight actions that deviated from the original action (e.g., incomplete actions, tilting the object, performing the actions with inefficient movement trajectories, etc.; 48 catch trial videos in total). All 240 videos were identical in terms of action timing, i.e., the videos started with hands on the table, followed by the action, and ended with hands moving to the same position of the table. The videos were edited in iMovie (Apple) and Matlab (MathWorks). Edited videos were gray scale, had a length of 2 s (30 frames per second), and a resolution of 400 × 225 pixels.

In the scanner, stimuli were back-projected onto a screen (60 Hz frame rate, 1024 × 768 pixels screen resolution) via a liquid crystal projector (OC EMP 7900, Epson Nagano, Japan) and viewed through a mirror mounted on the head coil (video presentation 6.9° × 3.9° visual angle). Stimulus presentation, response collection, and synchronization with the scanner were controlled with ASF (Schwarzbach, 2011) and the Matlab Psychtoolbox-3 for Windows (Brainard, 1997).

#### 2.1.3 fMRI Design

Stimuli were presented in an event-related design. In each trial, videos (2 s) were followed by a 1 s fixation period. 18 trials were shown per block. Each of the nine conditions (eight action conditions plus one catch trial condition) was presented twice per block. Six blocks were presented per run, separated by 10 s fixation periods. Each run started with a 10 s fixation period and ended with a 16 s fixation period. In each run, the order of conditions was pseudorandomized such that each condition followed each other condition equally often (max. once per block, on average 1.26 times per run). Each participant was scanned in a single session consisting of 8 functional scans and one anatomical scan. For each of the nine conditions there was a total of 2 (trials per block) × 6 (blocks per run) × 8 (runs per session) = 96 trials per condition. Each of the 24 exemplars per action condition was presented four times in the experiment.

#### 2.1.4 Task

Participants were instructed to attentively watch the movies. They were asked to press a button with the right index finger on a response button box whenever an action was meaningless or performed incompletely or incorrectly (i.e., in catch trials). Participants could respond either during the movie or during the fixation phase after the movie. To ensure that participants followed the instructions correctly, they completed a practice block outside the scanner.

#### 2.1.5 Representational dissimilarity matrices

In the initial analysis (GLM 1), we investigated five models that capture sociality, action direction, third-person-presence, movement trajectories, and object types present in the observed scenes (Fig. 3A).

For the Sociality model, participants judged the degree of sociality of the actions seen in the experiment. This rating was done after the fMRI session. For each of the eight actions, the participants were presented with two example frames of the beginning and the end of the action. The participants were asked to answer the following question on a 6-point Likert scale (from 1 = not at all to 6 = very much): “How much does the acting person takes into account the feelings and reactions of another person?” Outliers were defined as scores that deviated more than 2 standard deviations from the group mean (N=2 for *give cup* and *give cake*; N=1 for *place without social context*, *place without another person*, and *place onto saucer*). The resulting scores are shown in Table 1. The Sociality RDM was computed by averaging the rating scores across participants and subtracting each mean rating value from each other (Euclidean distance). We used rating averages rather than individual ratings to keep the orthogonality of the models included in the multiple regression (and thus the risk of potential collinearity) constant across participants. In an exploratory analysis, we also performed the multiple regression RSA with individual sociality models. The results of this analysis did not substantially differ from the results obtained by group-based sociality model and are therefore not discussed further.

**Table 1.** Rating results for sociality in Experiment 1.

|  | mean | SD | min | max |
| --- | --- | --- | --- | --- |
| give cup | 5.50 | 0.54 | 4.50 | 6.00 |
| place w/o soc | 2.13 | 1.07 | 1.00 | 5.00 |
| place w/o pers | 2.13 | 1.10 | 1.00 | 4.00 |
| place next | 3.23 | 1.15 | 1.00 | 5.50 |
| place saucer | 2.66 | 1.32 | 1.00 | 5.50 |
| take cake | 3.08 | 1.28 | 1.00 | 5.50 |
| take w/o pers | 1.93 | 1.10 | 1.00 | 4.00 |
| give cake | 5.67 | 0.51 | 4.50 | 6.00 |

The Third-Person-Presence RDM was computed by pairwise comparing each of the eight action conditions with each other with respect to whether they both contained a another person or not (0 = similar) or one condition contained another person and the other condition did not (1 = dissimilar).

For the Action Direction model, we defined four directions: toward the other person (irrespective of whether facing or not facing the actor), toward a specific target object (i.e., a saucer), toward no entity (unmarked location on the table), and toward the acting person his- or herself. The Action Direction RDM was computed by pairwise comparing each of the eight action conditions with each other with respect to whether they had the same action direction (0 = similar) or not (1 = dissimilar).

For the Movement Trajectory model, we defined three trajectories: movements away from the actor’s body in the frontal direction, movements away from the actor’s body to the left (place next to person), movements toward the actor’s body. The Movement Trajectory RDM was computed by pairwise comparing each of the eight action conditions with each other with respect to whether they had the same movement trajectory (0 = similar) or not (1 = dissimilar). Note that since we collapsed the stimuli across ‘actor side’ (left/right), this model did not capture information of absolute movement direction (e.g. from left to right; cf. the egocentric direction model of Experiment 2).

For the Object Type model, we defined three objects that were involved in the actions: a cup, a small plate/saucer, and a cake. The Object Type RDM was computed by comparing each of the eight action conditions with each other with respect to how many objects the actions share with each other (0,1, or 2).

In a second analysis (GLM 2), we specifically tested for sensitivity to discriminate between facing and nonfacing persons in a scene. To this end we constrained the analysis to the five action conditions that contain two persons (*give cup, place without social context*, *place next to person*, *take cake, give cake*). A model of Person Orientation was created by pairwise comparing each of the five action conditions with each other with respect to whether two action conditions both showed either facing or nonfacing persons (0 = similar) or whether one condition showed facing persons and the other condition showed nonfacing persons (1 = dissimilar; Supplementary Fig. 1A). Additional control models (Sociality, Action Direction, Movement Trajectory, Object Type) were created as described above (using the five action conditions containing two persons) and included in the regression.

In a third analysis (GLM 3), we investigated the four action directions in more detail. To this end we created dummy variables for each direction. RDMs were computed by pairwise comparing each action with each other with respect to whether both actions are directed toward the respective target or not (Supplementary Fig. 6A).

For each multiple regression, we tested for multicollinearity between the models by computing condition indices (CI), variance inflation factors (VIF), and variance decomposition proportions (VDP) using the colldiag function for Matlab. The results of these tests (GLM 1: CI<3, VIF<1.6, DVP<0.9; GLM 2: CI<4, VIF<3.3, VDP<0.9; GLM 3: CI<3, VIF<1.5, VDP<0.6) revealed no indications of potential estimation problems (Belsley et al., 1980).

#### 2.1.6 Data acquisition

Functional and structural data were collected using a 4 T Bruker MedSpec Biospin MR scanner and an 8-channel birdcage head coil. Functional images were acquired with a T2*-weighted gradient echo-planar imaging (EPI) sequence with fat suppression. Acquisition parameters were a repetition time of 2.2 s, an echo time of 33 ms, a flip angle of 75°, a field of view of 192 mm, a matrix size of 64 × 64, and a voxel resolution of 3 × 3 × 3 mm. We used 31 slices, acquired in ascending interleaved order, with a thickness of 3 mm and 15 % gap (0.45 mm). Slices were tilted to run parallel to the superior temporal sulcus. In each functional run, 176 images were acquired. Before each run we performed an additional scan to measure the point-spread function (PSF) of the acquired sequence to correct the distortion expected with high-field imaging (Zaitsev et al., 2004).

Structural T1-weigthed images were acquired with an MPRAGE sequence (176 sagittal slices, TR = 2.7 s, inversion time = 1020 ms, FA = 7°, 256 × 224 mm FOV, 1 × 1 × 1 mm resolution).

#### 2.1.7 Preprocessing

Data were analyzed using BrainVoyager QX 2.84 (BrainInnovation) in combination with the BVQXTools/NeuroElf Toolbox and custom software written in Matlab (MathWorks).

Distortions in geometry and intensity in the echo-planar images were corrected on the basis of the PSF data acquired before each EPI scan (Zeng and Constable, 2002). The first 4 volumes were removed to avoid T1 saturation. The first volume of the first run was aligned to the high-resolution anatomy (6 parameters). Data were 3D motion corrected (trilinear interpolation, with the first volume of the first run of each participant as reference), followed by slice time correction and high-pass filtering (cutoff frequency of 3 cycles per run). Runs, in which motion correction failed, were excluded (which was not the case for any run of any participant); apart from this criterion, no other criteria for exclusion due to motion artifacts was used. To assess head motion, relative (framewise) and absolute (runwise) head displacements were computed (Power et al., 2014). In no participant or run did mean framewise displacements exceed 0.57 mm (subject mean 0.25 ± 0.11 SD). Absolute displacements did not exceed 2.9 mm root mean square (rms) in any participant or run (subject mean 0.82 ± 0.37 SD). Spatial smoothing was applied with a Gaussian kernel of 8 mm FWHM for univariate analysis and 3 mm FWHM for MVPA (Wurm and Lingnau, 2015). For group analysis, both anatomical and functional data were transformed into Talairach space using trilinear interpolation. Univariate baseline contrasts (actions vs. baseline) and quality assessment for MVPA can be found in Supplementary Figures 11 and 12.

#### 2.1.8 Representational similarity analysis (RSA)

RSA (Kriegeskorte et al., 2008) was carried out using the CoSMoMVPA toolbox (Oosterhof et al., 2016a) and the representational similarity toolbox (Nili et al., 2014). For each participant and run, beta weights of the experimental conditions were estimated using design matrices containing predictors of the 8 action conditions, catch trials, and of 6 parameters resulting from 3D motion correction (x, y, z translation and rotation). Each predictor was convolved with a dual-gamma hemodynamic impulse response function (Friston et al., 1998). Each trial was modeled as an epoch lasting from video onset to offset (2 s). The resulting reference time courses were used to fit the signal time courses of each voxel. The resulting beta weights were averaged across the 8 runs to obtain one beta weight per condition and voxel. All searchlight (Kriegeskorte et al., 2006) and ROI MVPA were performed in volume space using spherical ROI with a radius of 12 mm.

For the ROI RSA, ROIs in left and right dorsal and ventral LOTC were defined based on the peak coordinates of the LOTC clusters identified in the searchlight decoding for sociality (Tal x/y/z, left DLOTC: −47/−58/4, right DLOTC: 48/−54/8) and transitivity (left VLOTC: −40/−61/−9, right VLOTC: 46/−55/−5), respectively, as reported in Wurm et al. (2017). Note that VLOTC ROIs were tested only in some additional analyses reported in Supplementary Materials. For each participant, ROI, and condition, we extracted and vectorized the beta values of the ROI to obtain one vector of beta values per action. For each vector, we demeaned the beta values across voxels by subtracting the mean beta value from each individual beta value. Next, we correlated the vectors with each other resulting in an 8 × 8 correlation matrix per ROI and participant. The neural correlation matrices were converted into a neural RDMs (1 – *r*). The lower triangulars of neural and model RDMs were vectorized, z-scored, and entered as independent and dependent variables, respectively, into a multiple regression RSA. Resulting beta coefficients were entered into one-sided signed-rank tests across participants (Nili et al., 2014). Statistical results were corrected for the number of ROIs and tested models (e.g., 2 ROIs * 5 models = 10 tests) using the false discovery rate (FDR) at q = 0.05 (Benjamini and Yekutieli, 2001).

Searchlight RSA was performed using identical parameters as reported above. For all searchlight analyses, individual beta coefficient maps were Fisher transformed and entered into one-sample *t*-tests (Oosterhof et al., 2016b). Statistical maps were corrected for multiple comparisons using an initial voxelwise threshold of *p* = 0.001 and 10000 Monte Carlo simulations as implemented in the CoSMoMVPA toolbox (Oosterhof et al., 2016a). Resulting z maps were used to threshold statistical maps (at *p* = 0.05 at the cluster level), which were projected on a cortex-based aligned group surface for visualization.

#### 2.1.9 Univariate fMRI analysis

Univariate activation effects were analyzed using univariate multiple regression. For each of the four action directions, we created dummy variables that encode for each action whether it is directed toward the respective target or not. In addition, we included a model encoding whether an action shows another person or not. As inclusion of all four action directions would result into perfect collinearity (“dummy variable trap”), we performed the multiple regression in a leave-one-model-out manner, i.e. we excluded each of the models in four iterative regressions. Resulting beta coefficients were averaged for each model and participant.

### 2.2 Experiment 2

#### 2.2.1 Participants

Twenty healthy adults (12 females; mean age, 24.7 years; age range, 19-32 years) not tested in Experiment 1 volunteered to participate in Experiment 2. All participants were right-handed with normal or corrected-to-normal vision and no history of neurological or psychiatric disease. One participant was excluded from the sample due to poor behavioral performance in the task (accuracy two standard deviations below the group mean). Participants gave written informed consent prior to participation in the study. The experimental procedures were approved by the Ethics Committee for research involving human participants at the University of Trento, Italy.

#### 2.2.2 Stimuli and Representational dissimilarity matrices

The goal of Experiment 2 was to investigate representations of implied person-directedness that are independent of whether the recipient is a third person present in the action scene or the observer her- or himself. To realize the perceptual cues that indicate observer-directedness while at the same time controlling as much as possible perceptual differences between the observer-directed actions and the actions directed toward other targets, we filmed actions taking place at a rectangular table from the viewpoint of a person sitting at this table (Fig. 2A). The acting person, another third person, and a target object were positioned at the remaining three sides of the table. The actions consisted of moving a grasped object into the four possible directions (observer, third person, object, actor). The target objects matched the respective moved objects (phone - charger, pen - pen holder, orange - fruit bowl). The videos started with the object already being grasped (hands in the middle of the table for the actor-directed actions; approx. 20 cm in front of the actor for the remaining action directions) and ended shortly before the reaching the target location, without making contact between objects and person or object targets. Videos had a length of 0.6 s. Only the upper torsos and arms of the actor and the third person were visible to move the observer’s viewpoint to the action scene as close as possible and to avoid unnecessary facial cues of the actor and the third person. For each action direction, 4 actions were used, resulting in 16 action conditions. In 8 of the 16 actions, a third person was shown to orthogonalize of a third-person-presence and person-directedness. Thus, in 50% of the person-directed actions (in all of the third-person-directed actions and in none of observer-directed actions), a third person was shown. To minimize covariance between implied person-directedness and various perceptual factors typically associated with the different action directions, we permuted (100000 iterations) all possible combinations of actor, third person, and object positions, third-person-presence, and action directions, and selected the 16 combinations that together result in the weakest correlations between the similarity model for implied person- directedness and control similarity models of various perceptual features of the actions. Specifically, the following control models were generated: (1) actor position (left, middle, right), (2) object position (left, middle, right), (3) third person position (left, middle, right, not present), (4) egocentric action direction (action directed toward the left, middle, right position from the observer’s viewpoint), (5) allocentric action direction (action directed toward the left, middle, right position from the actor’s viewpoint), (6) third-person-presence (present, not present), and (7) the distance between the end position of the hand/object and the observer’s viewpoint (close to observer (position 4 in Fig. 2A), middle of the table (positions 1 and 3 in Fig. 4A), far from observer (position 2 in Fig. 2A)). In addition, we generated models for each of the four action directions as well as an action direction model, similar to the direction models tested in Experiment 2 (Fig. 2C). For each iteration, the implied person-directedness model was correlated with the control models and the highest correlation coefficient was determined (“max”). From the 100000 iterations, the iteration with the lowest “max” correlation was selected (r(implied person-directedness – egocentric direction) = 0.12). Based on the combinations of positions and directions used in this iteration, the 16 action conditions were constructed. In addition to testing the correlation between implied person-directedness and control models of the final selection, we ensured that the other models do not show problematic correlations between each other by testing for collinearity between models used in the multiple regressions (GLM 1: CI<4, VIF<2.3, VDP<0.8; GLM 2: CI<4, VIF<2.5, VDP<0.8).

**Figure 4.**
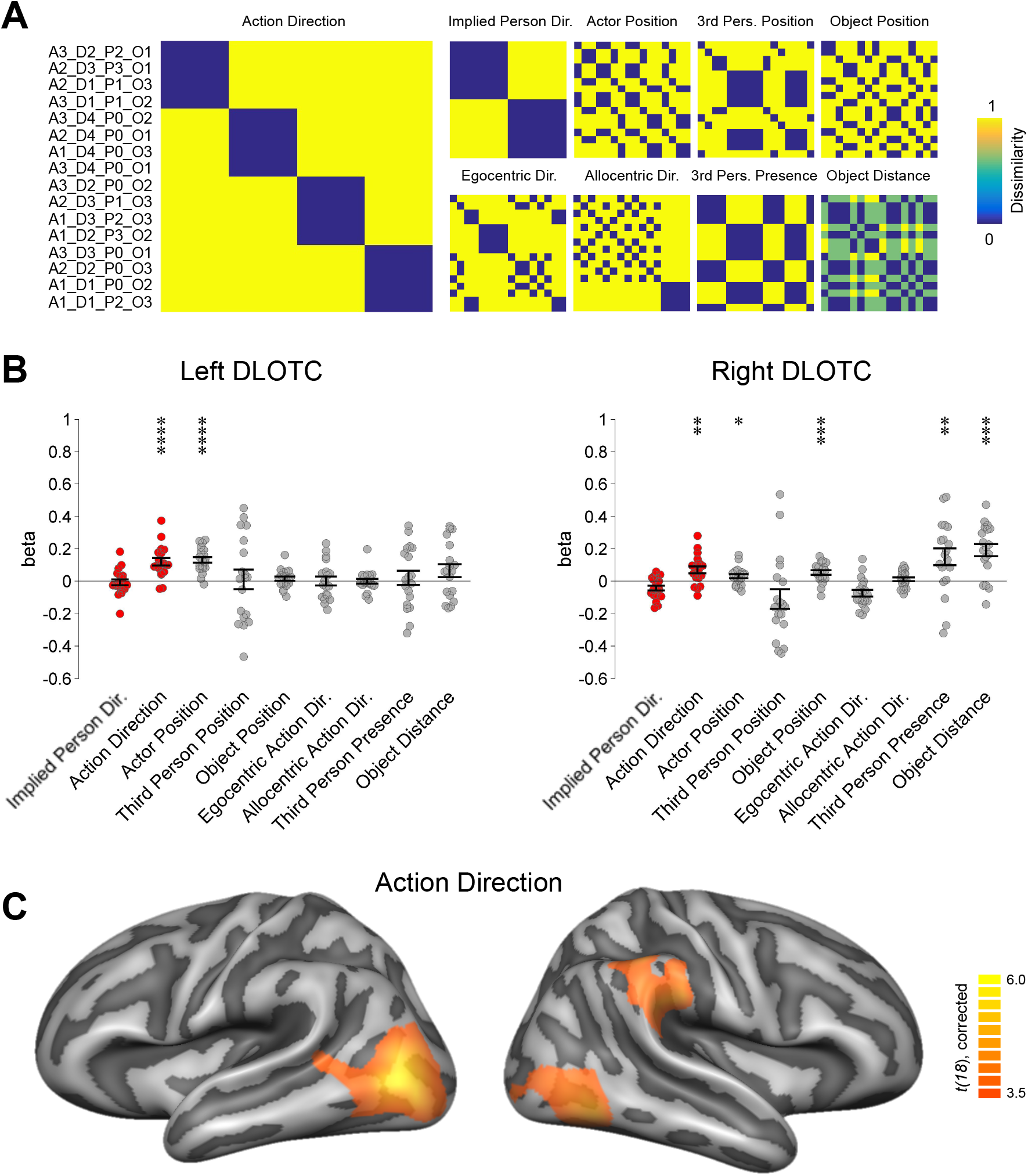
Multiple regression RSA targeting effects of person directedness and action direction (Experiment 2). (A) RDMs of tested models. (B) ROI multiple regression RSA. (C) Searchlight multiple regression RSA. Same conventions as in Figure 3.

To increase the perceptual variance within each action condition, each of the 16 actions were filmed in two different rooms, were performed with either the left or the right hand, and were performed with three types of objects (phone, pen, orange), resulting in 12 exemplars per condition (192 action videos in total; Fig. 2B).

Catch trials consisted of three exemplars of each of the 16 actions (48 catch trial videos in total). In these trials, the action direction changed in the middle of the video (e.g. in the first half of the video, it appeared that the action was directed toward the object, but in the second half of the video, the trajectory of the action was altered so that the action was ultimately directed toward the observer). All possible combinations from initial and end directions were used (e.g. if the original action was directed toward the object, then the catch trial versions started with the same direction but ended with observer, third person, or actor directions. The same video editing parameters and presentation settings were used as in Experiment 1.

#### 2.2.3 fMRI Design

Stimuli were presented in an event-related design. In each trial, videos (0.6 s) were followed by a 1 s fixation period. 36 trials were shown per block (each of the 16 action conditions were presented twice, plus 4 catch trials). Six blocks were presented per run, separated by 10 s fixation periods. Each run started with a 10 s fixation period and ended with a 16 s fixation period. In each run, the order of conditions was pseudorandomized such that each condition followed each other condition equally often (max. once per block, on average 0.35 times per run). Each participant was scanned in a single session consisting of 6 functional scans and one anatomical scan. For each of the 16 action conditions there was a total of 2 (trials per block) × 6 (blocks per run) × 6 (runs per session) = 72 trials per condition. Each of the 12 exemplars per action condition was presented 6 times in the experiment.

#### 2.2.4 Task

Participants were instructed to attentively watch the movies. They were asked to press a button with the right index finger on a response button box whenever an action changed its direction (i.e., in catch trials). Participants could respond either during the movie or during the fixation phase after the movie. To ensure that participants followed the instructions correctly, they completed a practice block outside the scanner.

To verify that observer-directed actions were perceived as directed toward the participants, they were asked after the fMRI session the following question: “There was one type of action in which an object was moved toward the camera viewpoint. How much did the action appear as if the object was given to you?” Participant responded using a Likert scale from 1 (“not at all; rather as if the object was moved toward a position on the table or given to someone not visible in the video”) to 6 (“very much; how it would feel like as if I was sitting at the table”). The mean response was 4.2 (SD = 1.1), suggesting that the observer-directed actions were sufficiently perceived as being directed toward the observing participants.

#### 2.2.5 Data acquisition, preprocessing, and analysis

The same neuroimaging and analysis procedures were used as in Experiment 1. In no participant or run did mean framewise displacements of head movements exceed 0.78 mm (subject mean 0.25 ± 0.18 SD). Absolute displacements did not exceed 4.0 mm root mean square (rms) in any participant or run (subject mean 0.66 ± 0.41 SD).”

#### 2.2.6 Behavioral control Experiment

Experiment 2 used shorter trial durations (0.6 s) than Experiment 2 (2 s). To test whether participants were able to differentiate the different action directions in Experiment 2, we conducted a behavioral control experiment that used the identical experimental design with the exception that no catch trials were included but participants responded to each trial at which target the action was directed at. Participants recognized the different action directions with high accuracy (recognition rate = 0.8 ± 0.09 SEM; chance = 0.25), and no significant differences in recognition rates and reaction times between the 4 directions were observed (recognition rates: F(3,15) = 1.23, p = 0.33; reaction times: F(3,15) = 0.8, p = 0.5; Supplementary Fig. 10). This suggests that the different action directions were equally well recognizable from the stimuli.

## 3. Results

### 3.1 Behavioral results

In Experiment 1, participants correctly detected catch trials with a rate of 96.2% (±1.0% SEM). In Experiment 2, participants detected catch trials with a rate of 88.4% (±2.2% SEM).

The generally high rate of correct responses suggests that participants attentively observed and recognized the actions. The lower detection rate in Experiment 2 was likely due to the shorter trial duration, which made the task more demanding.

### 3.2 Neuroimaging results

#### 3.2.1 Multiple regression RSA of Experiment 1

To characterize the representational content in LOTC in response to actions perceived as being social or nonsocial (Fig. 1), we computed a multiple regression RSA using a rating-based model of sociality, models of action direction, and third-person-presence as factors that are putative precursors of sociality, as well as control models for lower-level perceptual information (movement trajectory, object type; Fig. 3A). We specifically focused on characterizing the representational content of dorsal LOTC (DLOTC) we previously identified to be sensitive to the distinction of social vs. nonsocial actions (Wurm et al., 2017; for additional analyses in pSTS and in action observation and mentalizing networks, see Supplementary Materials). We found that the action direction model explained the representational dissimilarity in left DLOTC and third-person-presence explained the representational dissimilarity in both left and right DLOTC. By contrast, sociality as well as the other control models did not explain further variance (Fig. 3B).

A searchlight analysis using the same RSA procedure as in the ROI analysis revealed that the action direction model explained representational organization in left LOTC and lateral occipital cortex (LO; Fig. 3C, Tab. 2). Third-person-presence explained variance in bilateral LOTC, pSTS, and early visual cortex (EVC). The third-person-presence clusters in LOTC appeared to overlap with the extrastriate body area (EBA) (Downing et al., 2001): Peak coordinates of the clusters (Tab. 1) were close to coordinates obtained by functional localizers (e.g. Downing et al. (2001); Tal x/y/z, left EBA: −51/−71/11, right EBA: 52/−70/5). The effect of third-person-presence in EVC was likely due to the different amount of visual complexity in videos containing two persons as compared to one person. No brain area revealed significant effects for the sociality model and the control models (even when using more liberal correction thresholds).

**Table 2.** Clusters identified in Experiment 1. Coordinates (x, y, z) in Talairach space. Clusters corrected for multiple comparisons (voxel threshold p = 0.001, corrected cluster threshold p = 0.05), except for clusters marked with an asterisk (voxel threshold p = 0.001, uncorrected). Abbreviations: EVC, early visual cortex; LO, lateral occipital cortex; LOTC, lateral occipitotemporal cortex; pSTS, posterior superior temporal sulcus.

| Region | x | y | z | t | p |
| --- | --- | --- | --- | --- | --- |
| *Action Direction* |  |  |  |  |  |
| left LOTC | -42 | -64 | -2 | 7.75 | 2.68E-07 |
| left LO | -27 | -88 | 1 | 6.50 | 3.00E-06 |
| right LO | 24 | -88 | 4 | 6.02 | 9.00E-06 |
| *Person Presence* |  |  |  |  |  |
| left LOTC | -45 | -73 | 7 | 7.84 | 2.26E-07 |
| right LOTC | 45 | -67 | 4 | 6.58 | 3.00E-06 |
| left pSTS | -63 | -37 | 10 | 5.90 | 1.10E-05 |
| right pSTS | 51 | -34 | 13 | 4.79 | 1.28E-04 |
| EVC | -3 | -85 | 1 | 11.04 | 1.05E-09 |

We also explored whether LOTC is sensitive to the orientation of persons relative to each other in a scene. If so, representations of two facing persons (sitting at the same table) should be more similar with each other than with representations of two non-facing persons (sitting at different tables). Using multiple regression RSA restricted to the five action conditions that include two persons (Supplementary Figure 1A), we found that person orientation significantly explained the neural dissimilarity in left DLOTC over and above control models (Supplementary Figure 1B). A searchlight analysis demonstrated that the effect of person orientation was centered in left DLOTC dorsally and posteriorly of the clusters found for action directions and third-person-presence (Supplementary Figure 1C). A similar cluster at the same anatomical location was found in the right hemisphere, which however did not survive cluster size correction (Supplementary Table 1). In addition, we observed an effect in EVC (surviving correction in the right hemisphere only), which might be due to low-level visual differences between scenes with facing vs. non-facing persons.

Taken together, Experiment 1 revealed that basic perceptual action components that can be indicative of social actions (action directedness, third-person-presence, and person orientation), but not sociality as such, explain the representational content in LOTC.

#### 3.2.2 Multiple regression RSA of Experiment 2

The goal of Experiment 2 was to characterize representations of action directedness in more detail. Specifically, we aimed at testing whether the representation of person-directed actions requires the presence of another person or would also respond to actions *implied* to be directed toward other persons, such as actions directed toward the observer, who is typically part of an action scene. Conceptually, actions directed toward the observer are person-directed just like third-person-directed actions. But perceptually, the two types of person-directed action differ: Third-person-directed actions typically require the sight of another, third person in the scene whereas observer-directed actions do not. Additionally, the trajectories of observer-directed actions typically run along the observer-actor-axis, whereas the trajectories of third-person-directed actions typically point into different directions (unless the third person is positioned between the observer and the actor). There are hence two different types of directions – movements made in the direction of the observer and movements made into the direction of other persons – which could serve as cues for person-directedness. Does LOTC contain neural populations that generalize across these types of directions? Experiment 2 tested this question, using a design that allowed controlling the perceptual dimensions associated with the different kinds of action directions (third-person-, observer-, object-, and actor-directedness; Fig 2; see Methods for details).

Using multiple regression RSA, we tested a model of implied person-directedness against an action direction model as well as models of low- and mid-level perceptual dimensions that capture actor, person, and object positions, allocentric and egocentric action directions (i.e., to the left, across, or to the right from the actor’s or observer’s point of view, respectively), third-person presence, and the distance from the observer to the end position of the actor’s reach action (effectively modeling the size of the grasped object; Fig. 4A). In left DLOTC, the representational similarity of the observed actions could be explained by the action direction model and the actor position model (Fig. 4B). In right DLOTC, representational similarity could be explained by the action direction model and models for actor position, object position, third-person presence, and object distance. The implied person-directedness model did not explain variance over and above the action direction model.

A searchlight analysis revealed a cluster for the action direction model in left LOTC and additionally found clusters in right LOTC and right supramarginal gyrus (SMG; Fig. 4C, Tab. 3). The implied person-directedness model revealed no significant clusters (even when using more liberal correction thresholds). The control models for actor position and egocentric and allocentric action direction revealed effects in visual cortex, peaking in the occipital poles (Supplementary Fig. 2). Effects of actor position extended into LOTC bilaterally and overlapped with the cluster found for the action direction model (in left LOTC), but not with the cluster found for third person directedness (in right LOTC). The effect of third-person-presence extended into right, but not left LOTC and dorsally into SPL bilaterally. In contrast to Experiment 1, no effects of third-person-presence were observed in pSTS. The generally weaker effects of third-person-presence in higher-level visual areas in Experiment 2 are probably a consequence of the shorter trial duration (0.6 s as compared to 2 s in Experiment 1), which allowed for fewer saccades and thus less time to recognize the presence of another person, as well as the limited sight of the passive person, which likely reduced the salience of person-related information. Object position and object distance revealed clusters in right LOTC.

**Table 3.**
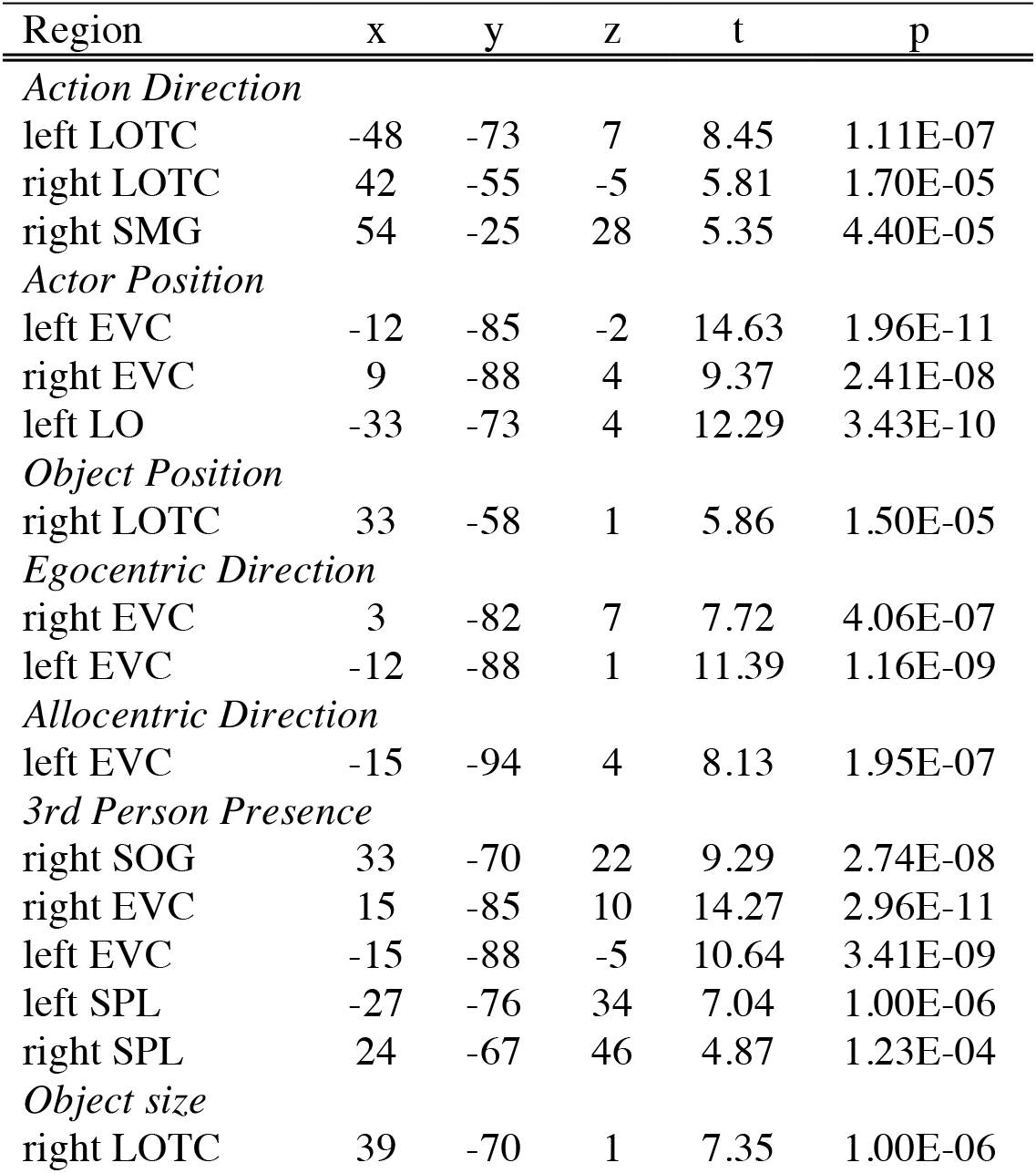
Clusters identified in Experiment 2. Coordinates (x, y, z) in Talairach space. Clusters corrected for multiple comparisons (voxel threshold p = 0.001, corrected cluster threshold p = 0.05). Abbreviations: EVC, early visual cortex; LO, lateral occipital cortex; LOTC, lateral occipitotemporal cortex; SMG, supramarginal gyrus; SOG, superior occipital gyrus; SPL, superior parietal lobe.

In conclusion, the multiple regression RSA of Experiment 2 replicated the effect of action direction in LOTC, and again revealed no evidence for a higher, interpretational level that represents implied person-directedness.

#### 3.2.3 Further analysis of action directedness

Both Experiment 1 and 2 revealed evidence that left LOTC encodes representations that capture the directedness of actions toward different types of recipients. This type of representation has not been described in the literature so far. We therefore explored representations of action directedness in more detail. First, we aimed at verifying that action direction effects in left LOTC were not affected by perceptual characteristics of movements associated with the different action directions. Notably, we included models that capture movement characteristics in both Experiment 1 (movement trajectories toward the actor or toward positions opposite or diagonally opposite of the actor) and 2 (movement directions from egocentric and allocentric points of view). To provide an additional control, we conducted a multiple regression RSA after including only actions with similar movement trajectories (Experiment 1: all actions except *take cake, take cake without other person, place next to person*; Experiment 2: third-person-directed and object-directed actions). This analysis revealed that effects of action direction were preserved in left LOTC (Supplementary Fig. 3), which corroborates the interpretation that the action direction effects observed in LOTC were independent of action movement information.

Second, we tested whether the identified representations specifically capture action directedness toward different types of entities (i.e., entity as *action recipient*) rather than the types of entities themselves (i.e., perceptual or conceptual information about persons or objects). For example, the entity targeted by an action might attract particular attention, which might result in increased neural processing of that type of entity. If so, we should observe increased neural activity in response to the respective targets, in particular for third-person-, actor-, and object-directed actions. By contrast, non-entity- and observer-directed actions are not directed toward a specific entity in the scene (a person or an object) and would therefore not be expected to trigger increased activity related to the processing of specific types of entities. To test these predictions, we analyzed univariate effects of the different action directions in the LOTC clusters identified in the searchlight multiple regression RSA. We used univariate multiple regression, i.e., the univariate responses to the conditions (Supplementary Figure 4A and B) were modeled with predictors for the different action directions and control variables (Experiment 1: third-person-directedness, object-directedness, non-entity-directedness, actor-directedness, and third-person-presence; Experiment 2: third-person-directedness, object-directedness, observer-directedness, actor-directedness, and third-person-presence; see Methods for details). In both experiments, action direction models could not explain the variance in activation in LOTC (Supplementary Figure 4C and D), except for actor-directed actions which revealed a negative effect, suggesting that activity was decreased for actor-directed actions. This effect might be explained by shorter movement trajectories, and thus less motion information, for actor-directed actions in Experiment 1. No other action direction model revealed significant effects (all p > 0.19). Taken together, these results do not match the pattern that would be expected if attention toward the different action targets resulted in differential activation of entity representations (i.e., increased neural responses to third-person-, actor-, and, object-directed actions but not to non-entity- and observer-directed actions).

Additional analyses, in which we explored the representational organization of action directions in more detail, are reported as supplementary materials (Supplementary Results, Supplementary Figures 5-7).

## 4. Discussion

The present study aimed at characterizing the representational content in LOTC in response to observed social and nonsocial actions. We focused on two different levels of description: (1) perceptual levels that capture socially relevant components of observed action scenes and (2) higher interpretational levels that capture the degree to which an action is implied to be social and person-directed. We found that areas in posterior LOTC encode representations that are sensitive to the directedness of actions toward different types of recipients (such as persons, objects, or the acting person herself) as well as other perceptual cues important for social interpretation (the presence of another person in the scene and the orientation of persons toward each other). By contrast, we found no evidence for representations that encode the perceived degree of sociality (Experiment 1) or representations of implied person-directedness (Experiment 2). These findings suggest that LOTC encodes salient perceptual components that are important for social interpretation rather than more abstract representations that capture interpretational descriptions of social actions. Notably, also in other areas downstream of LOTC, such as pSTS and areas of mentalizing and action observation networks, we did not find evidence for representations of sociality and implied person-directedness.

Our findings support a componential view of representational content in LOTC and argue against the hypothesis that LOTC encodes more complex and holistic representations of intentional aspects such as sociality. This interpretation fits well with the finding that LOTC is sensitive to different body parts (Bracci et al., 2015; Orlov et al., 2010; Taylor and Downing, 2011) and the interaction of body parts with external objects in the service of actions (Bracci and Peelen, 2013). In general, these representations might capture which functions are supported by a body part or hand-object interaction. Our main finding that LOTC is sensitive to the directedness of actions toward different types of targets provide evidence for an additional, more integrated level of representation that may constitute a critical intermediate processing stage during action recognition. How could action directedness be computed based on the perceptual information in an action scene? A straightforward solution could be that LOTC integrates information about the movement direction of a body part with categorical information about objects that lie within the area targeted by the body part. LOTC seems to be well situated to accomplish this type of integration: The action directedness cluster was centered in the middle occipital gyrus and closely overlapped with functional areas responsive to body parts (Downing et al., 2001), basic motion (Tootell et al., 1995), as well as action-related body motion (Beauchamp et al., 2002). These functional subregions thus might provide information about body and effector orientation and motion direction. Additionally, LOTC contains subregions that preferentially respond to different object categories, most prominently animate entities and inanimate manipulable entities (Chao et al., 1999; Konkle and Caramazza, 2013). Taken together, action directedness representations in LOTC might integrate movement and body information (where is a body part or movement directed at) with knowledge about entities (e.g. about animacy or manipulability). This stage of representation might serve as the perceptual basis for the interpretation of social actions and for the access of conceptual representations of actions in lateral posterior temporal cortex (Wurm and Caramazza, 2019). Notably, the observed sensitivity to discriminate different action directions cannot be explained by movement characteristics of the different actions (e.g. body movements like arm flexion for actor-directed actions vs. arm extension for action directed at other targets or movement trajectories to the left vs. to the right): First, we explicitly controlled for movement characteristics using models that captured type and direction of body movements. Second, action directions could still be discriminated after including only those actions with identical body movements. In addition, sensitivity to action directedness cannot be explained by increased processing of the recipient (e.g. the other person or the targeted object): Following this assumption, an action made in the direction of a particular target might direct attention toward that target, which would result in increased processing and accordingly increased activation of person- and/or object-responsive neural populations. The analysis of univariate neural responses did not find evidence for this interpretation. In conclusion, left LOTC appears to encode representations that discriminate different types of action recipients rather than types of entities per se.

LOTC was also sensitive to other components important for the recognition and interpretation of (social) actions, such as the presence of other persons and the orientation of persons relative to each other. While the finding of person orientation was exploratory and requires further investigation, it is notable that this type of representation was located in a different subregion of LOTC, i.e., posteriorly and dorsally of the action directedness cluster and the EBA and in proximity to the occipital place area (OPA), which responds to spatial aspects of scenes and scene elements (Dilks et al., 2011; Kamps et al., 2016). This area therefore seems to be suited to integrate information about body parts and body posture with spatial information about persons and objects in a scene. The view that the visual system contains representations sensitive to the orientation of persons relative to each other in a scene is also supported by the behavioral finding that facing persons are recognized more accurately then non-facing persons (Papeo et al., 2017). Taken together, dorsal posterior LOTC seems to capture facing persons, perhaps to group them into a single functional unit and to thereby provide an additional critical precursor for the detection of potential social interactions.

In contrast to the positive evidence for different action components in LOTC, both experiments revealed – after carefully controlling perceptual factors – no evidence for more abstract representations that capture more subjective levels of action interpretation, i.e., whether an action is implied to be social and person-directed. The absence of evidence is generally difficult to interpret. In our study, this is particularly critical because no other brain region showed effects of sociality. It therefore remains speculative whether the absence of effects was due to experimental aspects or points toward the non-existence of sociality representations. However, since both Experiment 1 and 2 failed to demonstrate significant – or at least trending – sociality effects, we think that our results might invite rethinking how abstract and complex intention-related aspects like sociality could be represented in the brain. Instead of a single type of representation that is specialized to capture sociality as such, there might be more distributed networks of brain areas that encode different aspects of actions (some of which we located in the present study) and that together provide the information for interpreting actions in a flexible and situation-dependent way. Following this view, the interpretation of an action in terms of sociality (and perhaps other abstract dimensions like morality, selfishness, etc.) does not occur in a single brain area but rather is formed by the interplay of different areas, perhaps under guidance of prefrontal areas that might integrate different sources of information depending on the specific situation and the focus of the observer. This view is supported by studies that consistently found activation of medial and ventrolateral prefrontal cortex when making different types of judgments about social actions (e.g. in terms of social status, morality, or intention), which might point toward a role of these areas in higher level integration and interpretation (Mason et al., 2014; Sinke et al., 2010; Wang et al., 2015). In addition, these studies found diverging effects in occipital and temporal areas, which might point toward differential involvement of these areas in processing different perceptual action aspects depending on a given task and available information. Manipulating task conditions, and thus the processing goal of the perceiver (Quadflieg & Koldewyn, 2017), is a necessary research program to investigate more thoroughly how access of higher level social dimensions in frontal areas and more basic socially relevant perceptual dimensions in posterior areas changes as a function of task, and which aspects are spontaneously extracted from stimuli (cf. Hafri, Trueswell, & Strickland, 2018). Note that the proposed interpretation does not rule out that other areas in the proximity (or part) of the visual system, such as pSTS and posterior middle temporal gyrus (pMTG), represent more general and complex aspects of actions in a task- and stimulus-independent manner. For example, areas in pSTS respond not only during the perception of social interaction but also to a variety of stimuli that could be socially relevant (e.g. facial expressions, gaze, vocal sounds, body movements, etc.), suggesting that pSTS has a role in reading out and integrating aspects about persons and actions that are potentially important in social situations, perhaps in the service of mental state inference (Deen et al., 2015; Lahnakoski et al., 2012; Yang et al., 2015). Likewise, action representations (e.g. of opening, giving, scratching, and agreeing) in pMTG generalize across observed action scenes and written sentences and are organized following semantic principles, suggesting that posterior temporal cortex represents actions at a conceptual level (Wurm and Caramazza, 2019).

Notably, the view that social interpretations do not build on a single unitary type of representation remains speculative until explicitly tested and we cannot exclude the possibility that the absence of sociality effects was due to stimulus- or task-related aspects. For example, by focusing on the minimal conditions sufficient for a social interpretation, the social cues might have been too subtle to (sufficiently strongly) activate representations sensitive to sociality. Another noteworthy aspect is that by manipulating various socially relevant cues in Experiment 1, it is possible that these cues might conflict with each other. For example, conflicts in interpretation might occur when persons face each other but the action is directed next to the person or when persons do not face each other but the action is directed toward the person. It is unclear how such conflicts might affect the activation of representations of social dimensions. Finally, our definition of sociality captured a general, high-level aspect of social action. Our results therefore cannot be generalized to other definitions capturing related dimensions such as mutual interactivity. These issues could be tested by expanding the range of factors that can be indicative of social actions (such as perceived and implied mutual interactivity, as in e.g. *communicating in person vs. via phone*, eye contact, etc.) and testing for their interactions.

On a final note, our results and interpretation seem to disagree with recent proposals that a subregion in right pSTS is selectively involved in the processing of social interactions (Isik et al., 2017; Walbrin et al., 2018). While there are methodological objections that concern the identification of social-interaction-responsive pSTS (see Introduction), both studies used MVPA to demonstrate that pSTS encodes representations that reliably discriminate between different kinds of social actions (animations of geometric shapes helping vs. hindering each other). However, it is not clear whether discrimination was driven by features specific for social interactions or lower-level perceptual features and whether pSTS would also discriminate between two comparably complex nonsocial actions. Therefore, besides theoretical counterarguments mentioned in the Introduction, the available results are too ambiguous to draw strong conclusions about the role of pSTS in processing exclusively social interactions. The apparent discrepancy of the results in pSTS could be resolved by the interpretation that pSTS responds to aspects that covaried with mutual interactivity in Isik et al. and Walbrin et al. but not in the present study, such as contingency between acting agents (Gao, Scholl, & McCarthy, 2012; Georgescu et al., 2014). In our study, there were no mutual interactions or reactions of the passive person, which might explain why no differential effects were observed in pSTS.

In summary, we found that areas in posterior LOTC are sensitive to basic action components that are critical for social interpretation and action recognition in general, but not to the perceived degree of sociality and implied person-directedness. These findings may help clarify which types of representations are encoded in LOTC and raise important questions of how more complex, interpretational levels of action description might be represented in the brain. We propose that interpretational descriptions of sociality and other abstract social dimensions are computed more flexibly in a situation- and task-dependent manner, by integrating variable types of information under control of downstream areas.

## 5. Acknowledgements

This work was supported by the German Research Foundation (DFG Research Grant WU 767/1-1), the Provincia Autonoma di Trento, and the Fondazione CARITRO (SMC). We thank Valentina Brentari for assistance in data acquisition and Paola Menapace and Giacomo Ariani for assistance in preparing stimulus material.

## Data and Code Availability Statement

Behavioral data, raw (dicom) and preprocessed (univariate glm) neuroimaging data, and the tested models of Experiment 1 and 2 are deposited at the Open Science Framework (https://osf.io/7ju9y/). Stimulus materials and code are available upon reasonable request.

## Notes

https://osf.io/7ju9y/

