## Supplementary Materials for "Lateral occipitotemporal cortex encodes perceptual components of social actions rather than abstract representations of sociality"

### Supplementary Material

**Supplementary Results**

To explore representational organization of action directions tested in the experiments in more detail, we performed additional analyses. Note that the in-depth analysis of action directions is narrowed by two limitations: First, in Experiment 1, there are fewer actions per action direction and the number of actions is not equal for each action direction (e.g. there were 3 third-person-directed actions and only one object-direct action), which might complicate the identification of action-direction-specific clusters. Second, only three action-directions were common across both experiments (third-person-, object-, and actor-directed actions), whereas non-entity- and observer-directed actions occurred only in Experiment 1 and 2, respectively, which limits the comparability of studies.

We first investigated the representational organization of action directions relative to each other using multidimensional scaling (MDS). The inspection of MDS revealed that the different action directions were separated in representational space without salient subdistinctions between (or superordinate clustering of) the different action directions (Supplementary Figure 5).

We then explored whether LOTC contains representations that selectively distinguish one action direction from the remaining action directions. To this end, we created four separate models for the different action directions (Experiment 1: third-person-, object-, actor-directedness, and directedness toward no specified entity; Supplementary Figure 6A; Experiment 2: third-person-, observer-, object-, and actor-directedness; Supplementary Figure 7A). ROI analyses were performed in DLOTC as described in the main article. In addition, we included the ventral LOTC (VLOTC), which revealed a distinction between object-directed and not object-directed actions (Wurm et al., 2017; see also Wurm & Caramazza, 2019).

The ROI multiple regression RSA for Experiment 1 revealed that only third-person- and object-directedness explained variance in left and right LOTC (Supplementary Figure 6B). Effects of third-person-directedness were significant in left and right DLOTC, whereas effects of object-directedness were significant in left VLOTC and, less strongly, in left DLOTC. A repeated measures ANOVA with the factors MODEL (third-person-, object-directedness), ROI (DLOTC, VLOTC), and HEMISPHERE (left, right) revealed an interaction of MODEL x ROI ($F(1,19) = 7.16$, $p = 0.015$), which replicated the double dissociation of person- and object-directedness in DLOTC and VLOTC, respectively, in Wurm et al. (2017). In addition, we observed a main effect of hemisphere ($F(1,19) = 15.11$, $p = 0.001$), indicating that the models generally explained more variance in the left than in the right hemisphere. A searchlight analysis corroborated these findings: the third-person-directedness model explained variance in left DLOTC; the object-directedness model explained variance in left VLOTC, partially overlapping with the third-person-directedness cluster in DLOTC (Supplementary Figure 6C, Supplementary Table 1). The third-person-directedness model also revealed a cluster in right DLOTC, which however, did not survive correction for multiple comparisons. Both left and right DLOTC clusters revealed almost perfect overlap with the clusters found in Wurm et al. (2017) (3D peak distances left DLOTC: 2.4 mm, right DLOTC: 4.4 mm).

The ROI multiple regression RSA for Experiment 2 revealed that object-directedness, but not third-person-directedness, explained representational similarity of the actions in left DLOTC (Supplementary Figure 7B). The same trend was found in left VLOTC, but the effects were not significant. In right DLOTC, we observed an effect of third-person directedness and a weaker effect of object-directedness. In right VLOTC, we found the opposite pattern, i.e., a stronger effect of third-person-directedness and no effect of object-directedness. A repeated measures ANOVA revealed that these unexpected interactions between models and hemispheres as well as between models and ROIs were significant: In left LOTC, representational organization was best explained by object-directedness whereas in right LOTC, representational organization was best explained by third-person-directedness (interaction of MODEL x HEMISPHERE: ($F(1,18) = 37.27$, $p < 0.001$). Object-directedness explained representational organization better in dorsal as compared to ventral LOTC, whereas third-person-directedness explained representational organization better in ventral as compared to dorsal LOTC (interaction of MODEL x ROI: ($F(1,18) = 8.32$, $p = 0.01$). Paired samples *t*-tests revealed no significant differences between dorsal and ventral ROIs, for both left and right ROIs and both third-person- and object-directedness (all $p > 0.08$). A searchlight analysis corroborated these observations and revealed no additional effects in other brain areas (Supplementary Figure 7C, Supplementary Table 2). Observer-directedness revealed effects in bilateral (but more strongly left) SPL and left PMd, in line with our expectations. Actor-directedness revealed an effect in early visual cortex, which is likely due to differences in trajectory lengths of the reaching movements, which were shorter for the actor-directed as compared to the remaining actions. A similar but not significant effect of actor-directedness was found in Experiment 1.

In summary, Experiment 1 revealed representations of third-person- and object-directedness in dorsal and ventral LOTC, respectively, consistent with previous studies (Wurm et al., 2017; Wurm and Caramazza, 2019). Experiment 2 also identified representations of third-person- and object-directedness in LOTC, but no consistent anatomical overlap with Experiment 1 (Supplementary Fig. 4). Most strikingly, effects of third-person-directedness were found in right but not left LOTC. As there were several experimental differences between Experiment 1 and 2 with regard to trial duration (2 s vs. 0.6 s, respectively), stimulus details (whole person vs. hands and torso only, respectively), and inclusion of different action directions (non-entity-directed vs. observer-directed actions, respectively) it remains speculative what could be the reason for these anatomical inconsistencies. A possible explanation for the absence of third-person-directedness in left LOTC could be that also third-person-*presence* effects were lateralized to the right hemisphere, i.e., absent in left LOTC (Supplementary Figure 2). Following this interpretation, the visual third-person cues were too weak (possibly due to the shorter trial duration and limited sight of the person) to sufficiently trigger representations of third-person presence, a prerequisite of third-person-directedness, in left LOTC.

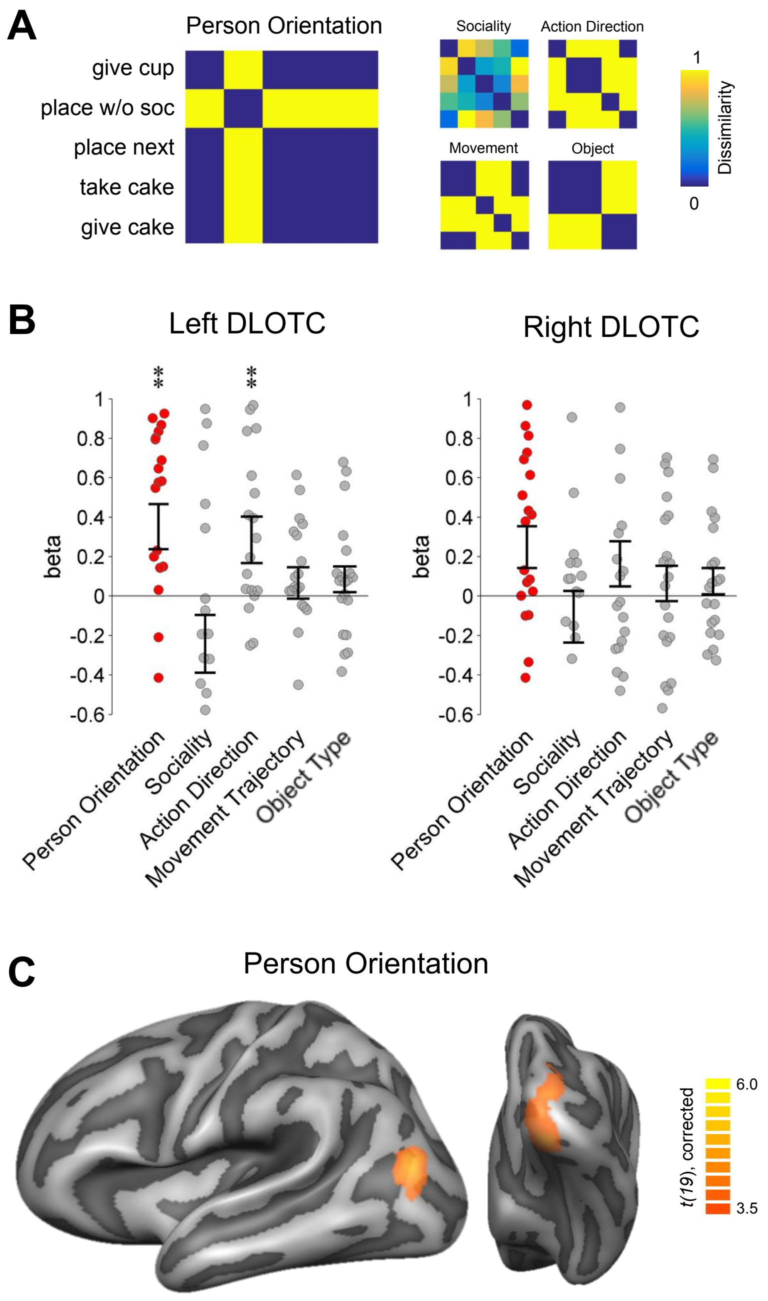


**Supplementary Figure 1.** RSA targeting effects of person orientation (Experiment 1, GLM 3). (A) RDMs of the tested models. (B) ROI multiple regression RSA in DLOTC. No significant effects were observed in VLOTC. A repeated measures ANOVA with the factors ROI (DLOTC, VLOTC), and HEMISPHERE (left, right) revealed a main effect of ROI ($F(1,19) = 12.95$, $p = 0.002$; no main effect of HEMISPHERE or interaction, both $p>0.2$), indicating that the effect of person orientation was restricted to the dorsal LOTC. Asterisks in indicate FDR-corrected (for number of models per test) effects: ** $p < 0.01$. Error bars indicate SEM. (C) Searchlight multiple regression RSA. Statistical maps are corrected for multiple comparisons (voxel threshold $p = 0.001$, corrected cluster threshold $p = 0.05$).

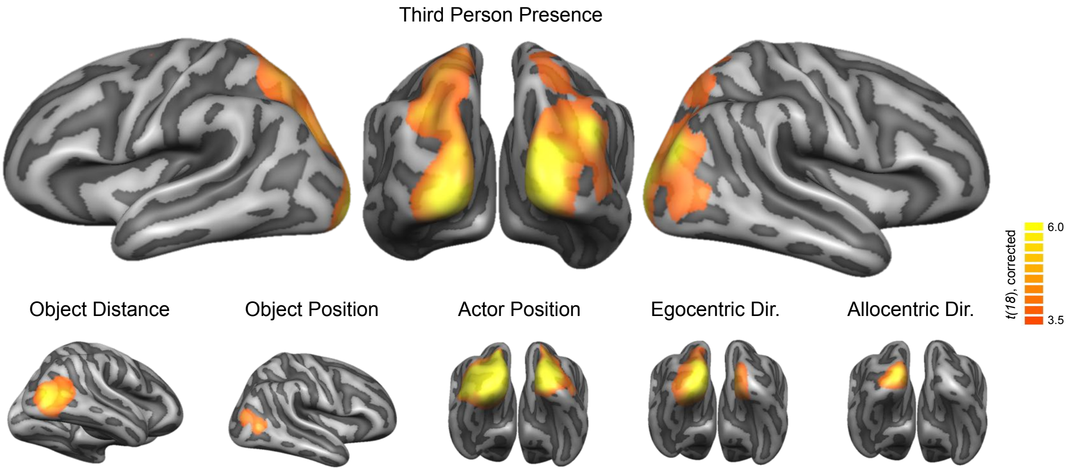


**Supplementary Figure 2.** Searchlight multiple regression RSA (Experiment 2, GLM 1; similar effects were revealed by GLM 2). Statistical maps are corrected for multiple comparisons (voxel threshold p = 0.001, corrected cluster threshold p = 0.05).

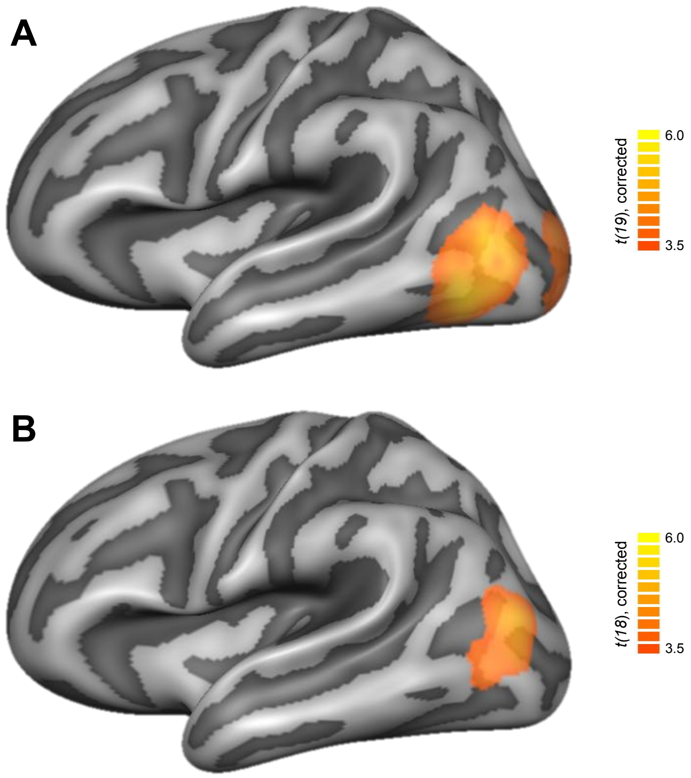


**Supplementary Figure 3.** Multiple regression RSA effects of the action direction model for actions with similar movement trajectories. (A) Experiment 1; after exclusion of self-directed actions (*take cake, take cake without other person*) and *place next to person*. (B) Experiment 2; after exclusion of observer-directed and self-directed actions. Statistical maps are corrected for multiple comparisons (voxel threshold p = 0.001, corrected cluster threshold p = 0.05).

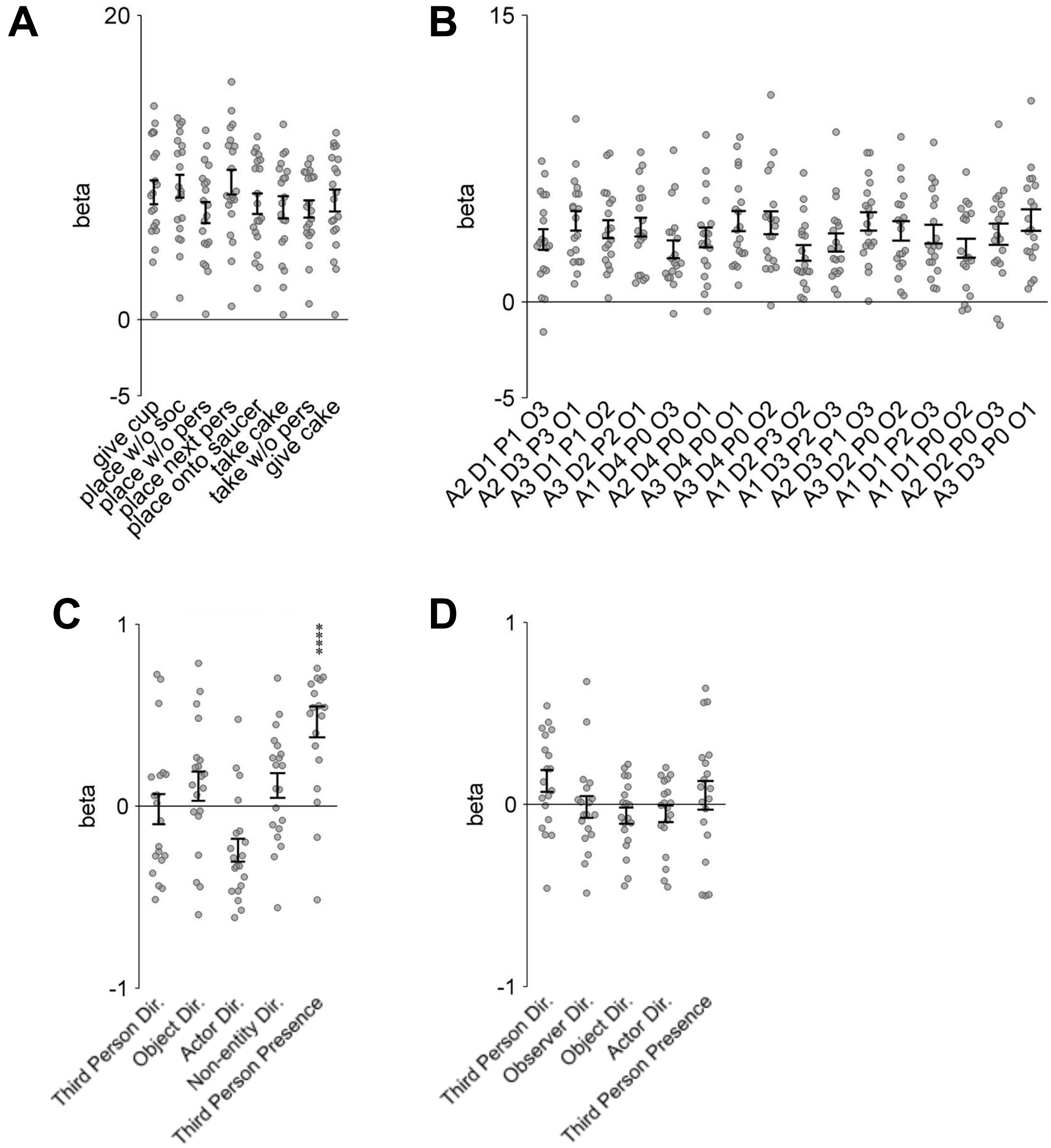


**Supplementary Figure 4.** Univariate multiple regression RSA. (A-B) Activation in response to each condition of Experiment 1 (A) and 2 (B). Beta coefficients of conditions were extracted from spherical (12 mm radius) around peak voxels revealed for the action direction model in the multiple regression RSA (Experiment 1: -42/-67/7, Experiment 2: -48/-73/7). (C-D) Univariate multiple regression for Experiment 1 (C) and 2 (D) using beta coefficients shown in A and B as dependent variables. Experiment 1 (C) revealed a significant effect of third-person-presence, which suggests increased processing demands in left LOTC during the observation of two as compared to one person (one-sided paired t-test, $t(19) = 5.47$, $p < 0.0001$). Actor-directed actions revealed a negative effect, suggesting that activity was decreased for actor-directed actions. This effect might be explained by shorter movement trajectories, and thus reduced motion information, for actor-directed actions in Experiment 1. Experiment 2 (D) revealed a trend for third-person-directedness, which however did not survive FDR correction (one-sided paired t-test, $t(18) = 2.14$, $p = 0.023$; all other $p > 0.18$). Error bars indicate SEM.

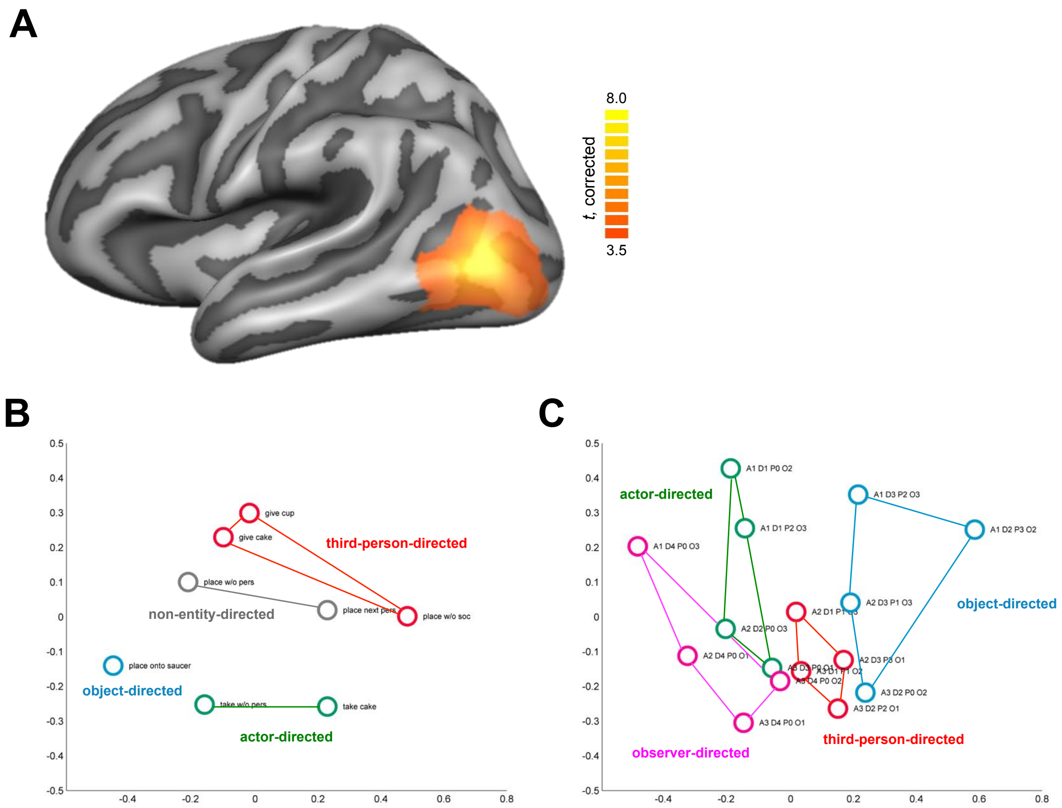


**Supplementary Figure 5.** Multidimensional scaling (MDS) analyses in the conjunction cluster of action directedness (A) for Experiment 1 (B) and 2 (C). MDS were performed on representational dissimilarity matrices (RDMs) using metric stress. RDMs were extracted from spherical ROIs (12 mm radius) around the peak voxel (Talairach coordinates, x/y/z: -42/-70/4) of the conjunction of statistical maps for the action direction model in the multiple regression RSA (voxel threshold p = 0.001, corrected cluster threshold p = 0.05).

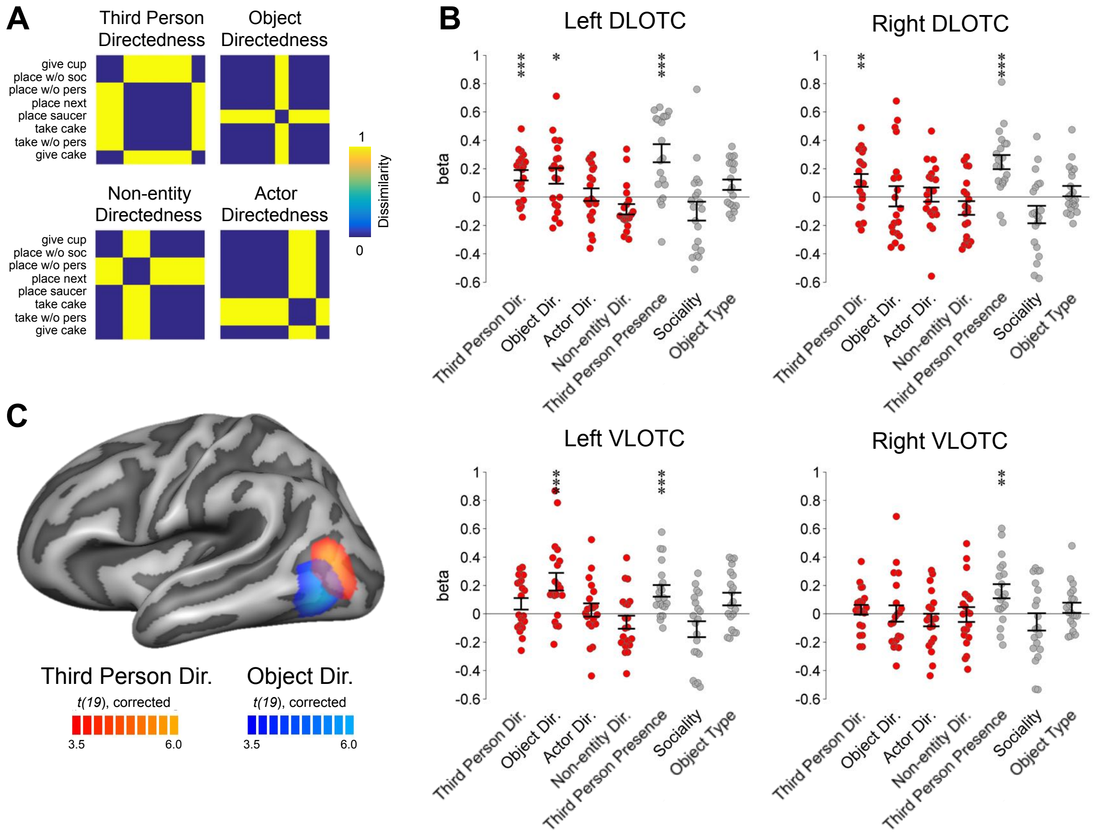


**Supplementary Figure 6.** RSA targeting effects of different action directions (Experiment 1, GLM 2). (A) RDMs of four action direction models. (B) ROI multiple regression RSA. Asterisks in indicate FDR-corrected (for number of models per test) effects: * $p < 0.05$, ** $p < 0.01$, *** $p < 0.001$, **** $p < 0.0001$. Error bars indicate SEM. (C) Searchlight multiple regression RSA. Statistical maps are corrected for multiple comparisons (voxel threshold $p = 0.001$, corrected cluster threshold $p = 0.05$).

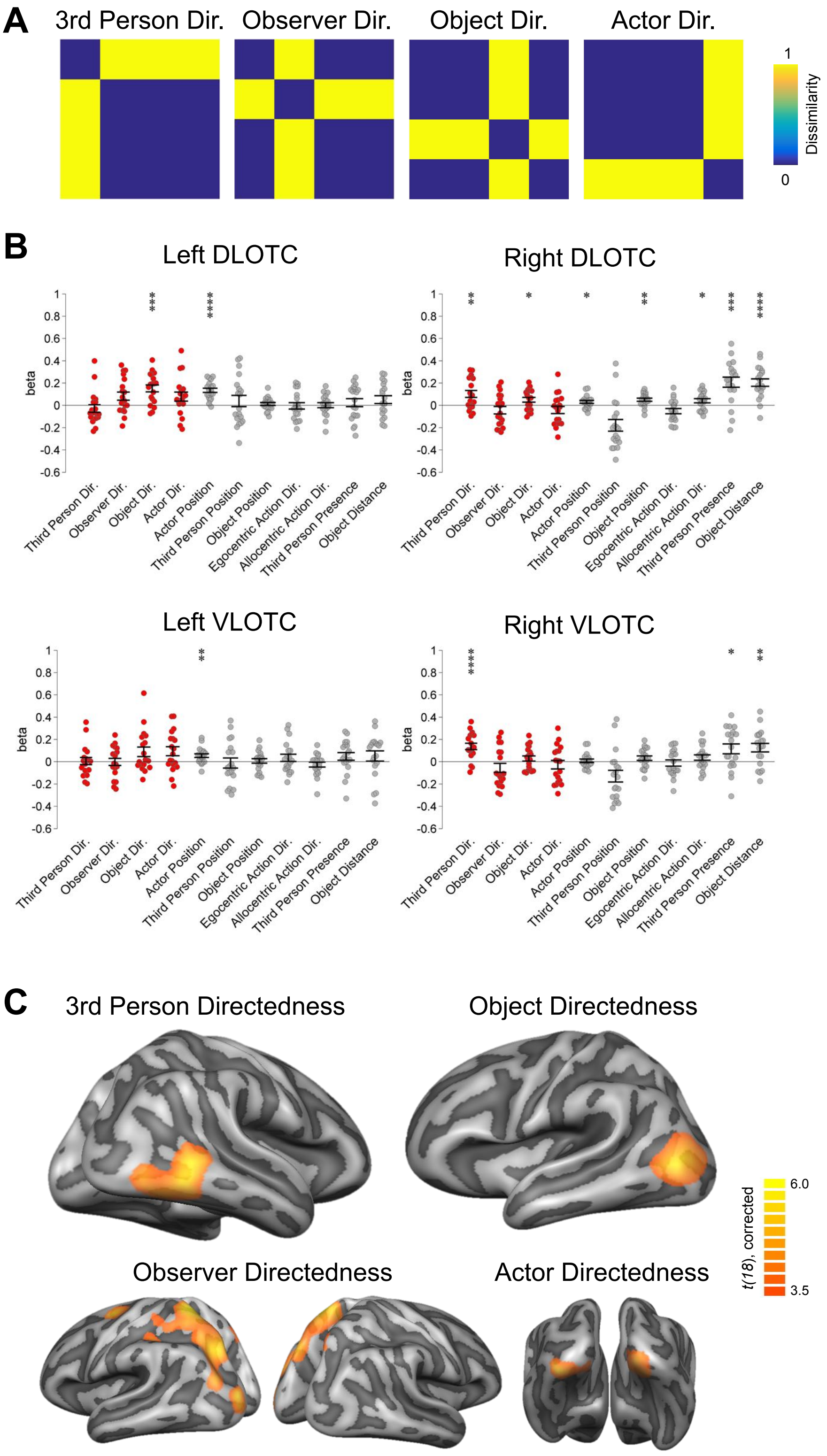


**Supplementary Figure 7.** RSA targeting effects of different action directions (Experiment 2, GLM 2). (A) RDMs of four action direction models. (B) ROI multiple regression RSA. Asterisks in indicate FDR-corrected (for number of models per test) effects: * $p < 0.05$, ** $p < 0.01$, *** $p < 0.001$, **** $p < 0.0001$. Error bars indicate SEM. (C) Searchlight multiple regression RSA. Statistical maps are corrected for multiple comparisons (voxel threshold $p = 0.001$, corrected cluster threshold $p = 0.05$).

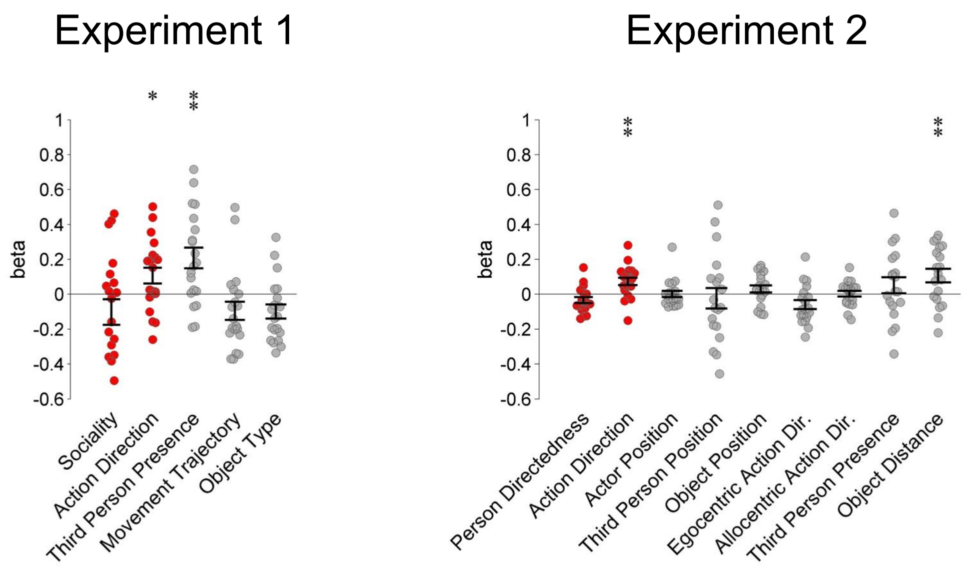


**Supplementary Figure 8.** ROI multiple regression RSA in right pSTS. ROIs (12 mm radius) were based on coordinates reported in Isik et al. (2017) for the contrast social interactions vs. two independent actions (Tal x/y/z, 53/-42/19; converted from MNI coordinates, 54/-43/18). Asterisks in indicate FDR-corrected (for number of models per test) effects: * $p < 0.05$, ** $p < 0.01$, *** $p < 0.001$, **** $p < 0.0001$. Error bars indicate SEM.

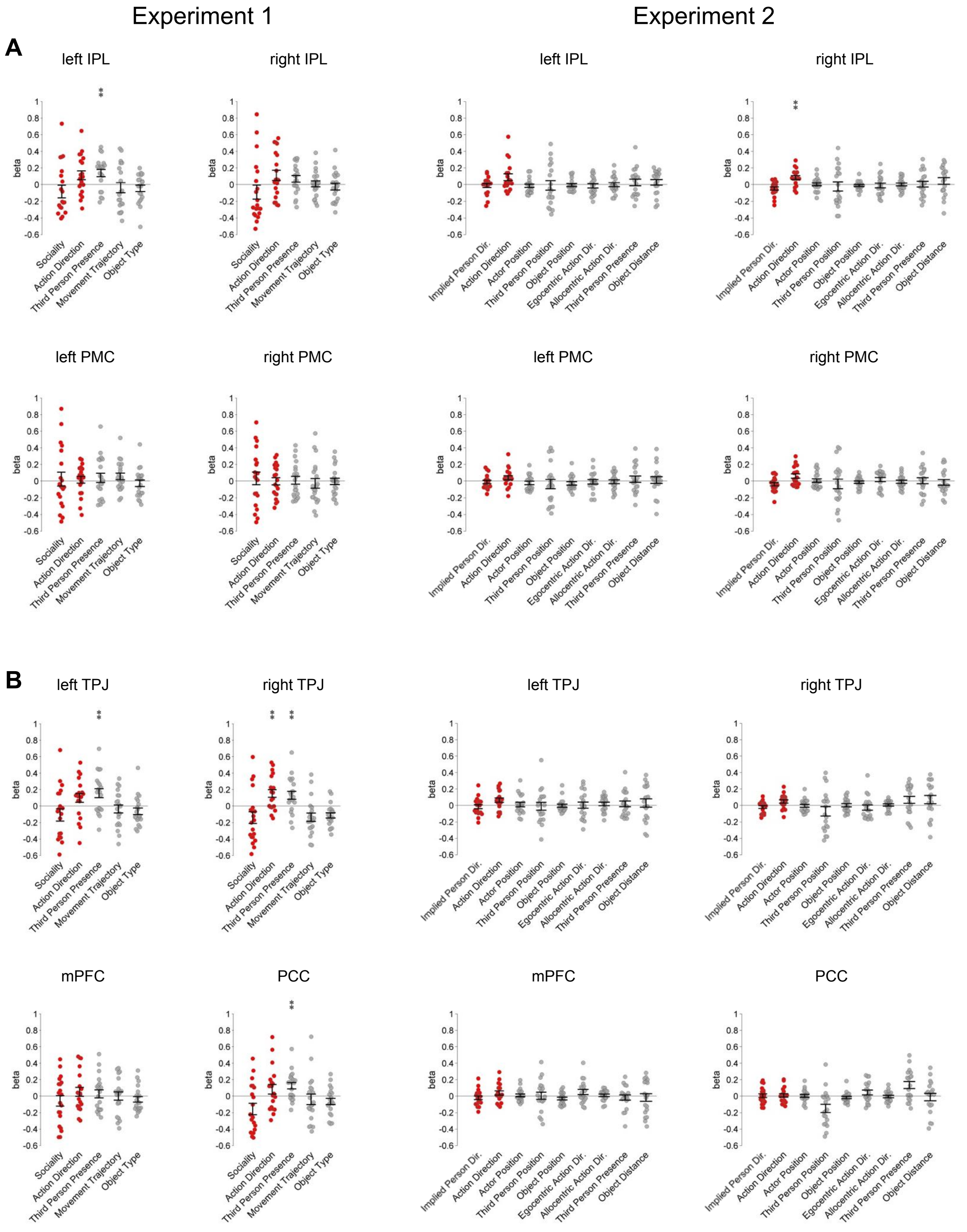


**Supplementary Figure 9.** ROI multiple regression RSA in frontoparietal areas of the action observation network (A) and the mentalizing network (B). ROIs (12 mm radius) were based on (A) peak coordinates reported in Caspers et al. (2010) for the contrast “action observation” (Tal x/y/z, left PMv: -50/9/30, right PMv: 52/12/26, left IPL: -60/-24/36, right IPL: 44/-34/44) and (B) coordinates as used in van Overwalle & Baetens (2009) for the mentalizing network (left TPJ: -50/-55/25, right TPJ: 50/-55/25, mPFC: 0/50/20, PPC: 0/-60/40). GLM 2 for action directions did not reveal any significant effects in both Experiments and is therefore excluded here). Asterisks in indicate FDR-corrected (for number of models per test) effects: * $p < 0.05$, ** $p < 0.01$, *** $p < 0.001$, **** $p < 0.0001$. Error bars indicate SEM.

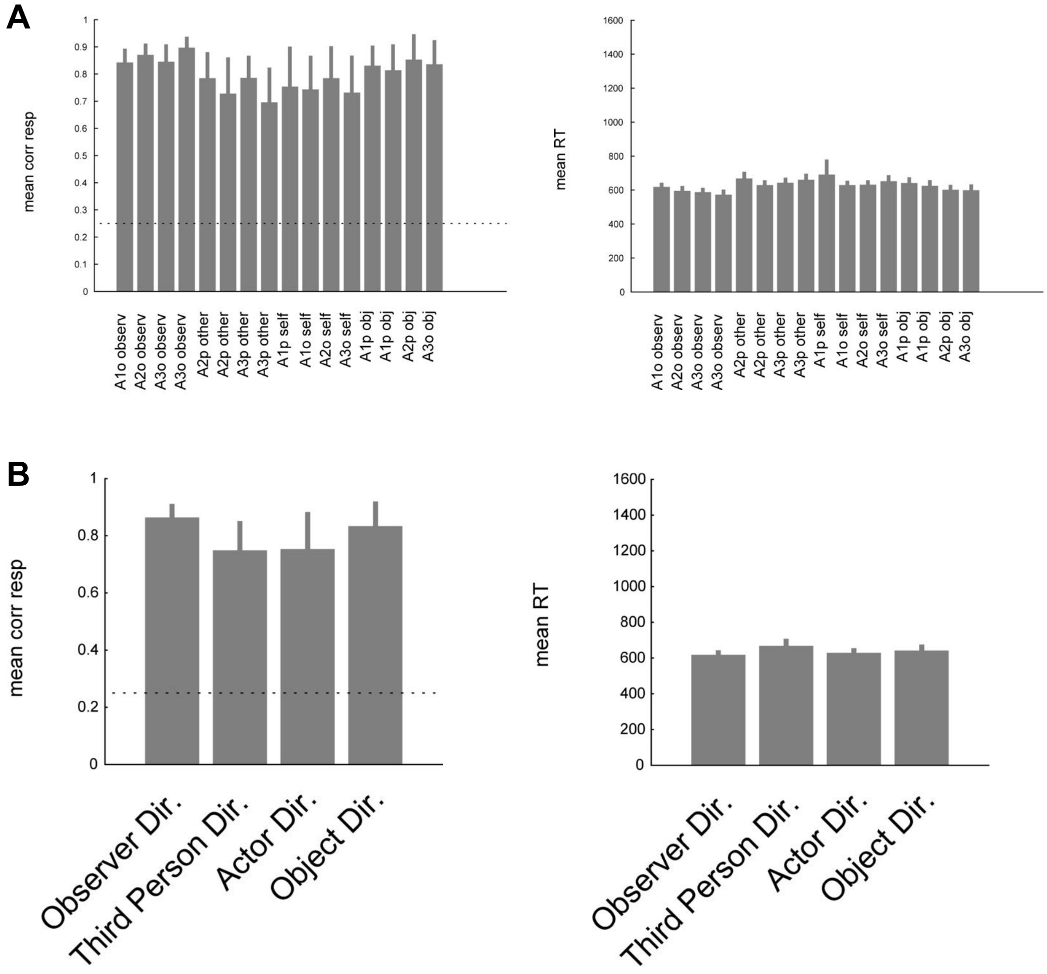


**Supplementary Figure 10.** Behavioral control experiment of Experiment 2. Experimental design was identical to Experiment 2 with the exception that only 3 runs were used, catch trials were excluded, and instead participants (N = 6) responded to each trial at which target the action was directed at. Responses were given by button press on a keyboard with the right hand (index to little finger, balanced across participants). (A) Recognition rates and reaction times to each of the 16 actions. (B) Recognition rates and reaction times to each of the four action directions collapsed across the 4 actions per direction. The dotted line indicates recognition rate at chance (0.25). Error bars indicate SEM.

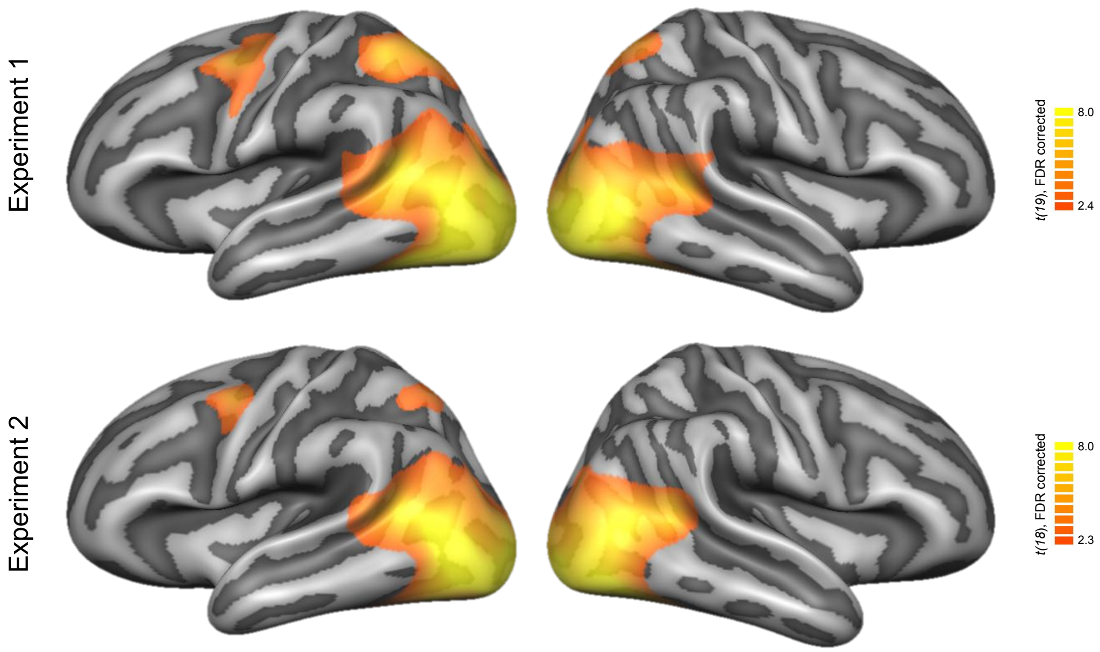


**Supplementary Figure 11.** Univariate RFX GLM maps for actions v. baseline.

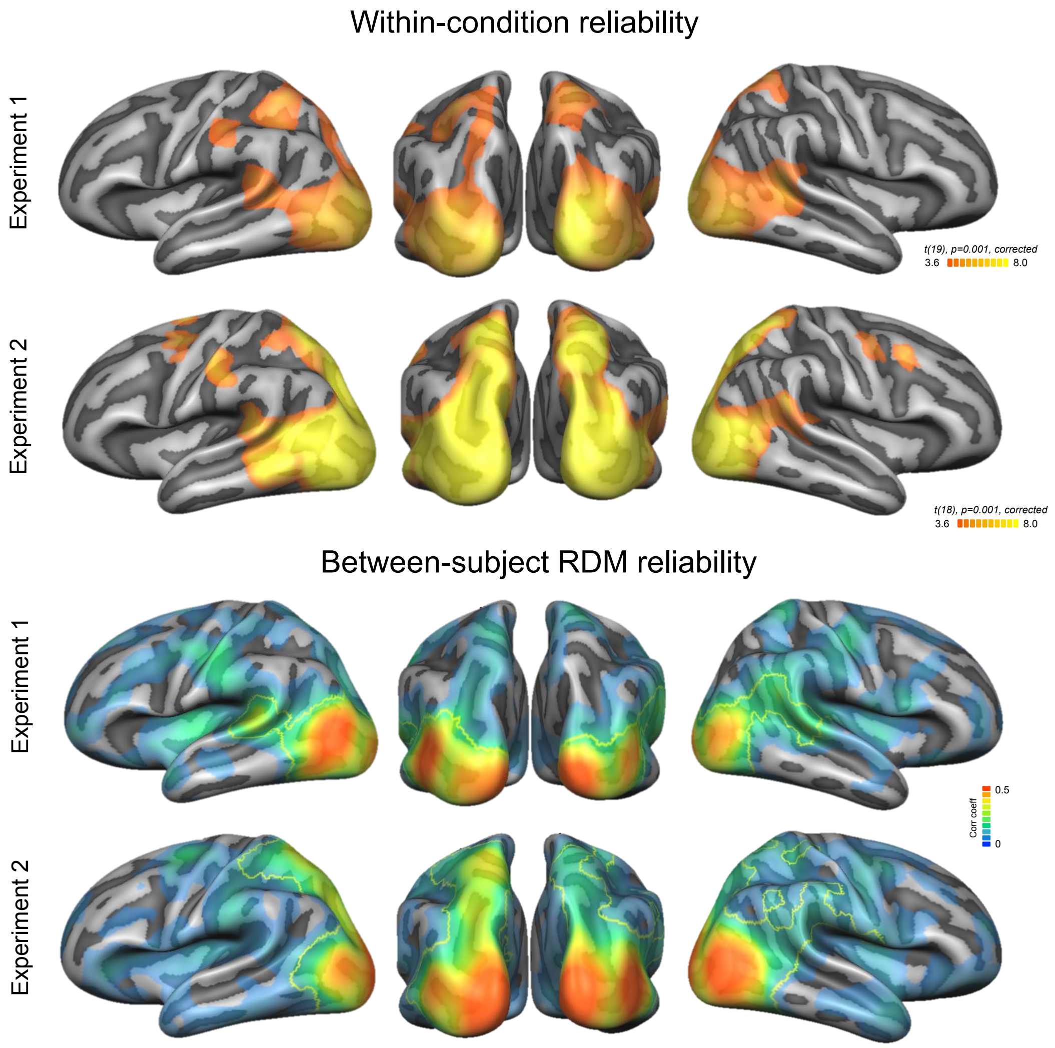


**Supplementary Figure 12.** Reliability of neural pattern data within and across participants. (A) To investigate whether neural patterns for the different conditions are reliably distinct from each other across runs, a searchlight correlation RSA was performed, i.e., for each searchlight sphere (12 mm), voxel patterns for each condition were correlated with each other using split-half cross validation (all possible combinations of runs) and the mean of off-diagonal cells of the correlation matrix was subtracted from the mean of the on-diagonal cells of the matrix (Ritchie et al., 2017). Differences were entered into paired samples t tests across participants, and t maps were thresholded using corrections for multiple comparisons (voxel threshold $p = 0.001$, corrected cluster threshold $p = 0.05$). (B) To investigate whether the neural dissimilarities between conditions are reliably similar across participants, a between-subject RSA was performed, i.e., for each searchlight sphere (12 mm), neural RDMs were computed (as described in Methods). For each participant and searchlight sphere, RDMs were correlated with the mean RDM of the remaining participants (leave-one-subject-out cross validation), and resulting correlation coefficients were entered into paired t tests. Maps show mean correlation coefficients, outlines indicate clusters corrected for multiple comparisons (voxel threshold $p = 0.001$, corrected cluster threshold $p = 0.05$). Resulting maps highlight areas in which the models tested in the Experiments should generally be able to explain representational content of the neural data.

Supplementary Table 1. Clusters identified for different action directions and person orientation in Experiment 1

| Region | x | y | z | t | p |
|---|---|---|---|---|---|
| *Person Directedness* | | | | | |
| left DLOTC | -39 | -67 | 7 | 5.52 | 2.50E-05 |
| *Object Directedness* | | | | | |
| left VLOTC | -48 | -61 | -11 | 4.45 | 2.76E-04 |
| *Person Orientation* | | | | | |
| left DLOTC | -42 | -73 | 13 | 5.04 | 7.30E-05 |
| right DLOTC* | 39 | -64 | 16 | 4.85 | 1.10E-04 |
| right EVC | 21 | -76 | -8 | 4.73 | 1.50E-04 |

Coordinates (x, y, z) in Talairach space. Clusters corrected for multiple comparisons (voxel threshold p = 0.001, corrected cluster threshold p = 0.05), except for clusters marked with an asterisk (voxel threshold p = 0.001, uncorrected). Abbreviations: EVC, early visual cortex; (D/V)LOTC, (dorsal/ventral) lateral occipitotemporal cortex.

Supplementary Table 2. Clusters identified for different action directions in Experiment 2

| Region | x | y | z | t | p |
|---|---|---|---|---|---|
| *Third Person Directedness* | | | | | |
| right LOTC | 51 | -55 | -5 | 5.49 | 3.30E-05 |
| *Observer Directedness* | | | | | |
| right SPL | 18 | -76 | 55 | 4.85 | 1.27E-04 |
| left SPL | -15 | -52 | 52 | 5.76 | 1.80E-05 |
| left PMd | -21 | -4 | 55 | 5.14 | 6.80E-05 |
| left TOS | -27 | -91 | 10 | 6.55 | 4.00E-06 |
| *Object Directedness* | | | | | |
| left LOTC | -39 | -70 | 4 | 7.73 | 3.98E-07 |
| *Actor Directedness* | | | | | |
| left EVC | -30 | -79 | -14 | 5.43 | 3.70E-05 |
| right EVC | 6 | -79 | -11 | 5.60 | 2.60E-05 |

Coordinates (x, y, z) in Talairach space. Clusters corrected for multiple comparisons (voxel threshold $p = 0.001$, corrected cluster threshold $p = 0.05$). Abbreviations: EVC, early visual cortex; LOTC, lateral occipitotemporal cortex; PMd, dorsal premotor cortex; SPL, superior parietal lobe; TOS, transverse occipital sulcus.
